# *polo* affects cell fate upon ionizing radiation in *Drosophila* hematopoietic progenitors by negatively regulating *lok*

**DOI:** 10.1101/2023.06.22.546047

**Authors:** Tram Thi Ngoc Nguyen, Guang-Chao Chen, Jiwon Shim, Young-Han Song

## Abstract

In response to ionizing radiation (IR), stem cells undergo cell cycle arrest, senescence, premature differentiation, or cell death. The decision between survival and death is critical during tumorigenesis and effective killing of cancer cells. We used the larval *Drosophila* lymph gland, a hematopoietic organ, as a model to understand the mechanism for cell fate decisions during stem cell development. The hematopoietic progenitors survived or died via apoptosis when larvae were irradiated in early or late third instar larval (L3) stages, respectively. In late L3 progenitors, the basal level of *polo* (*Drosophila* PLK1) was low, enabling IR-induced activation of *lok* (*Drosophila* CHK2), which was necessary and sufficient for inducing autophagy and reactive oxygen species (ROS) production resulting in cell death. The high level of *polo* in early L3 progenitors negatively regulated *lok* resulting in significantly low or undetectable levels of ROS or autophagy, respectively. The surviving early L3 progenitors underwent cell cycle arrest followed by premature differentiation affected by *tefu* (*Drosophila* ATM) and *lok* mutation. These results provide clues for designing effective therapeutic strategies for cancer.

**Summary statement:** We elucidated the mechanism underlying cell fate decisions during stem cell development in larval *Drosophila*, which will help develop effective cancer treatment modalities.

## Introduction

DNA damage induced by genotoxic agents, such as ionizing radiation (IR), activates the DNA damage response (DDR) in cells, resulting in cell cycle arrest, DNA repair, senescence, and programmed cell death. Besides these responses in somatic cells, stem cells undergo premature differentiation to bypass the harmful effects of accumulated DNA damage in the stem cell pool (Barazzuol et al., 2017; Inomata et al., 2009; Matsumura et al., 2016). The fate of cells with damaged DNA depends on the severity and nature of the DNA damage, genetic status, and cell type. Defective regulation of DDR in stem cells can lead to genomic instability and cancer. Various chemotherapeutic agents and radiation can induce DNA damage and kill cancer cells by activating the DDR pathway. Therefore, understanding the mechanism of cell fate decisions, especially between survival and death, can provide clues to understanding tumor formation and designing effective strategies to treat cancer.

In mammals, DNA damage-induced cell death occurs through ATM, CHK2, and p53 activation, inducing the expression of pro-apoptotic genes, PUMA and NOXA. Notably, p53, a major tumor suppressor gene, plays an important role in the decision between survival and death via various mechanisms (Liu et al., 2021). Besides causing DNA damage, DNA-damaging agents, including IR, induce reactive oxygen species (ROS) production (Rowe et al., 2008) and initiate macroautophagy (also called autophagy) (Chaurasia et al., 2016; Czarny et al., 2015)—a lysosome-mediated pathway that degrades and recycles damaged macromolecules and organelles (Das et al., 2012). Although ROS functions as a signaling molecule at low levels, excessive amounts of ROS lead to cell death (Azad et al., 2009). Autophagy is generally considered to be necessary for cell survival in both basal and induced states; however, it can result in cell death in certain cellular contexts (Doherty and Baehrecke, 2018). Additionally, ROS regulates autophagy through several mechanisms including upregulation of Beclin 1, oxidation of Atg4, and mitochondrial dysfunction, further complicating the signaling pathway (Redza-Dutordoir and Averill-Bates, 2021). However, the roles of ROS, autophagy, and their crosstalk in the DDR that determines cell fate decisions in stem cells remain unclear.

Hematopoiesis in *Drosophil*a occurs during larval development via the proliferation and differentiation of hematopoietic progenitors in the medullary zone (MZ) located in the primary lobe of the lymph gland (Banerjee et al., 2019). After the active proliferation phase during early larval stages, hematopoietic progenitors enter proliferative quiescence in the late third instar larval stage (L3) and differentiate into mature hemocytes, including plasmatocytes and crystal cells occupying the periphery of the lymph gland, known as the cortical zone (CZ). Additionally, lamellocytes, which are rarely detected in healthy larvae, are induced in the CZ under stress conditions, such as wasp infestation. We previously showed that apoptotic cell death occurs in hematopoietic progenitors after IR in the late L3 stage in a *tefu* (*Drosophila* ATM)-, *lok* (*Drosophila* CHK2)-, and *dp53* (*Drosophila* p53)-dependent manner (Nguyen et al., 2021). However, less is known about the DDR in hematopoietic progenitors during larval development, despite extensive studies on stress response in the hematopoietic system. For example, developmental signals are utilized to increase differentiation in response to stresses including hypoxia, nutrient deprivation, and wasp infestation (Banerjee et al., 2019; Shim, 2015). Moreover, during larval development, the basal level of ROS increases in the progenitors of the late L3 lymph gland, and ROS function as signaling molecules that induce differentiation (Owusu-Ansah and Banerjee, 2009). Nevertheless, to the best of our knowledge, the possibility that the high basal level of ROS in late L3 progenitors affects cell fate decision upon IR has not been determined.

This study aimed to investigate the DDR of hematopoietic progenitors during larval development and the mechanism underlying cell fate decision. IR induced premature differentiation or apoptotic cell death in progenitors irradiated during early or late L3 stages, respectively. Moreover, the expression level of *polo* (*Drosophila* Polo-like Kinase 1, PLK1) affected the cell fate decision by negatively regulating *lok*, which mediates IR-induced autophagy and ROS production, both of which are required for cell death. Since the level of PLK1, autophagy, and ROS are frequently altered in cancer cells, our results provide clues to understand tumor formation and may help toward designing effective strategies to treat cancer.

## Results

### DNA damage induced at the early L3 stage does not induce cell death in hematopoietic progenitors

Previously, we showed that hematopoietic progenitors in the lymph gland undergo apoptotic cell death when irradiated at L3 stage (88 h after egg hatching, AEH) (Nguyen et al., 2021). Since the cellular responses of stem cells vary depending on their developmental stages, we tested whether the apoptotic cell death of hematopoietic progenitors is induced in the earlier larval stages. When mid L3 (76 h AEH) or older larvae (80 or 84 h AEH, data not shown) were irradiated, a significant number of apoptotic cells marked by active cleaved *Drosophila* caspase (cDcp-1) was detected 4 h after irradiation in hematopoietic progenitors and differentiated cells (Fig. 1A, data not shown). In contrast, cell death was not detected when irradiated at early L3 stage (44 h AEH) (Fig. 1A). Furthermore, *I-CreI*-induced DNA double-strand breaks (DSBs), which increase cell death in the late L3 lymph gland (Nguyen et al., 2021), did not induce cell death in the early L3 lymph gland (Fig. S1A). Additionally, the hematopoietic niche called the posterior signaling center, marked by *Antp>GFP*, did not undergo IR-induced cell death when irradiated at both early and late L3 stages (Fig. S1B).

**Fig. 1.**
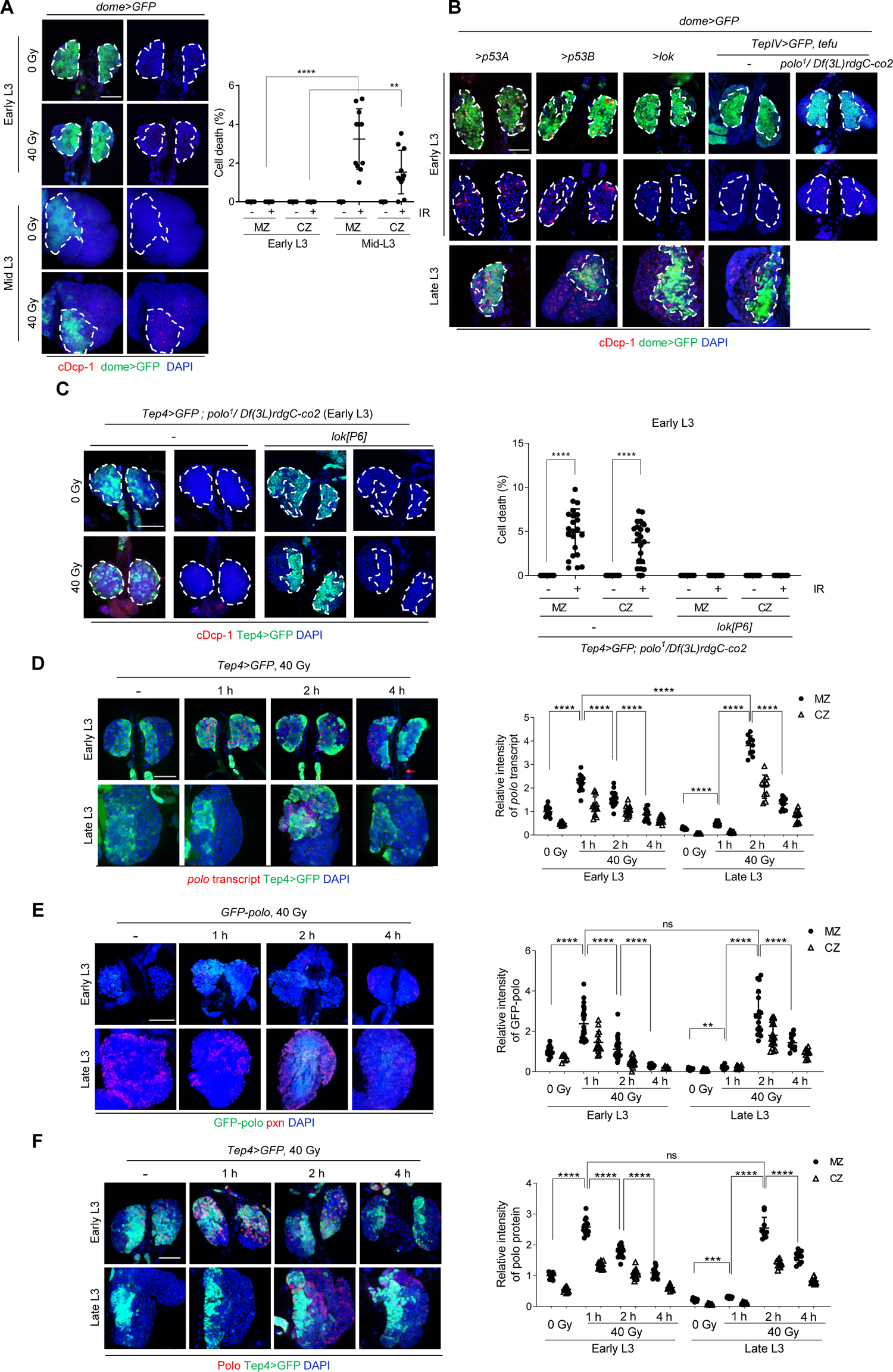
*lok* inactivation by a high level of *polo* transcripts mediates the survival of early L3 progenitors after irradiation. (A) The early and mid L3 larvae were irradiated at 40 Gy, and the lymph gland was stained with a cDcp-1 antibody to detect apoptotic cells 4 h after treatment. Representative images are shown. Scale bars, 50 μm. DAPI (blue), dome>GFP (green), and cDcp-1 (red) indicate DNA, progenitors, and apoptotic cells, respectively. The boundary of the dome>GFP-stained MZ is marked with white broken lines. Percentages of cell number with cDcp-1 signal in progenitors (MZ) and differentiated cells (CZ) are shown. Each data point represents a single primary lobe (n>11), and mean±s.d. are shown. **** *P<*0.0001, ** *P<*0.01. (B) The lymph glands from early or late L3 larvae that overexpress genes in the cell death pathway including *dp53A, dp53B, lok, or tefu* in wild type or *polo* mutant background in MZ by *dome-Gal4* were stained using a cDcp-1 antibody. DAPI (blue), dome>GFP (green), and cDcp-1 (red) indicate DNA, progenitors, and apoptotic cells, respectively. (C) IR-induced cell death was determined at 4 h after 40 Gy irradiation of early L3 larvae with indicated genotypes. DAPI (blue), Tep4>GFP (green), and cDcp-1 (red) indicate DNA, progenitors, and apoptotic cells, respectively. Percentages of cell number with cDcp-1 signal in progenitors (MZ) and differentiated cells (CZ) are shown. Each data point represents a single primary lobe (n>10), and mean±s.d. are shown. (D–F) The *polo* expression level was evaluated via RNA *in situ* hybridization using an anti-sense *polo* probe (D), GFP-POLO reporter (E), or anti-Polo antibody (F) in early or late L3 lymph gland at 0 Gy and 1, 2, and 4 h after 40 Gy irradiation. DAPI (blue), Tep4>GFP (green), Pxn (red), and GFP-POLO (green) indicate DNA, progenitors, differentiated cells, and GFP-polo fusion protein, respectively. The intensity of *polo* transcript (D), GFP-POLO (E), and Polo protein (F) in MZ and CZ marked by Tep4>GFP or Pxn are shown. The values are normalized relative to the polo transcript or GFP-POLO intensity in the early L3 MZ before irradiation. Each data point represents a single primary lobe (n>10), and mean±s.d. are shown. **** *P<*0.0001, *** *P<*0.001, ** *P<*0.01, not significant. The boundaries of MZ are marked with white broken lines. Scale bars, 50 μm.

### *lok* inactivation by *polo* mediates the survival of early L3 progenitors after irradiation

We have previously shown that IR-induced cell death in late L3 hematopoietic progenitors requires *tefu*, *lok*, and *dp53*, which induces the pro-apoptotic genes, *hid* and *reaper* (Nguyen et al., 2021). To investigate the survival mechanism of early L3 hematopoietic progenitors upon irradiation, these genes were overexpressed in the MZ using the *Gal4-UAS* system. Overexpression of the two major *dp53* isoforms (Zhang et al., 2014), *dp53A* and *dp53B*, or their upstream kinase, *lok*, using *dome-Gal4* induced cell death in the early and late L3 progenitors (Fig. 1B), suggesting that cell death pathway downstream of *lok* is functional in early L3 progenitors. Moreover, the *lok*-induced cell death required Lok kinase activity because the overexpression of kinase-dead Lok did not induce cell death (data not shown). Additionally, we used a misexpression line (*tefu^GS13617^*) (Gregory et al., 2007) containing the upstream activating sequence (UAS) recognized by GAL4 to determine the role of *tefu*, the upstream kinase of *lok*. When *tefu* was expressed with a single MZ population driver, either *Tep4-Gal4* or *dome-Gal4*, no cell death was detected (data not shown). However, when Tefu protein levels were upregulated using two Gal4 drivers, cell death was detected in the late L3 lymph gland (Fig. 1B), suggesting that a relatively high amount of *tefu* expression is required to induce cell death in the progenitor. In contrast, *Tep4*- and *dome-Gal4*-driven overexpression of *tefu* in early L3 progenitors was insufficient to induce cell death (Fig. 1B). These results suggest that the absence of IR-induced cell death in early L3 progenitors was because of the lack of *lok* activation upon irradiation.

CHK2 is a protein kinase activated by ATM-mediated phosphorylation in response to DNA damage (Zannini et al., 2014). Several mechanisms of CHK2 inactivation have been reported in mammals, including dephosphorylation by protein phosphatase 2A (Freeman et al., 2010), proteasomal degradation (Bohgaki et al., 2013), and inhibitory phosphorylation by PLK1 (Tsvetkov et al., 2005; van Vugt et al., 2010). We tested the possibility of negative regulation of *lok* by *polo* to understand the mechanism underlying the absence of *lok* activation in early L3 hematopoietic progenitors after irradiation. Since the *polo* null mutant is lethal, we used an allelic combination of a hypomorphic *polo^1^*and a deficiency line (*Df(3L)rdgC-co2*) with complete deletion of the whole region comprising the *polo* gene. In this *polo* mutant, at early L3 stage, many apoptotic cells were detected after irradiation (Fig. 1C). Moreover, the loss of *lok* restored the absence of IR-induced cell death in the early L3 *polo* mutant (Fig. 1C), suggesting that IR-induced activation of *lok* is blocked by *polo* in early L3 progenitors, resulting in survival after irradiation. Similar results were obtained when *polo* RNAi reduced *polo* levels in the MZ (Fig. S2A). Additionally, *polo* mutation restored the lack of cell death owing to *tefu* overexpression in early L3 stage (Fig. 1B). Furthermore, overexpression of the constitutively active form of *polo* (*UAS-poloT182D*) (Khaminets et al., 2020) using *dome-Gal4*, which did not affect cell death in the absence of irradiation, significantly attenuated IR-induced cell death in the *dome>GFP*-positive MZ (5.5%) compared with that in the control (11.4%) when irradiated at late L3 stage (Fig. S2B). As an internal control, IR-induced cell death in the *dome>GFP*-negative CZ—where *poloT182D* was not expressed—was unaffected (Fig. S2B). Additionally, *lok* overexpression-induced apoptotic cell death in larval eye discs was attenuated by co-expressing *poloT182D* (Fig. S2C). These results suggested that *polo* negatively regulates cell death by inactivating *lok*.

### *polo* expression level during the development of the lymph gland

We determined *polo* levels using fluorescence *in situ* hybridization (FISH) to understand the mechanism underlying developmental stage-specific regulation of *lok* in progenitors. Without irradiation, the percentage of *polo*-positive cells in MZ was 27.5-fold higher in early L3 stage (22%) than that in late L3 stage (0.8%). Moreover, *polo* transcripts in late L3 stage (0.279 ± 0.048) were almost undetectable, and the intensity of *polo* in the early L3 progenitors (0.999 ± 0.204) was 3.6-fold higher than that in late L3 progenitors (Fig. 1D). In response to irradiation, *polo* transcripts rapidly increased to their maximum level (2.226 ± 0.345) at 1 h and decreased to the basal level by 4 h in early L3 stage. In contrast, a low basal expression of *polo* in the late L3 stage reached the maximum level (3.812 ± 0.408) at 2 h after IR and decreased rapidly (Fig. 1D). To test whether the polo level was regulated at the transcriptional level, we used the *polo*-reporter*, GFP-POLO* (Moutinho-Santos et al., 1999). The expression patterns of *GFP-POLO* (Fig. 1E) and the *polo* transcript were similar. Moreover, the Polo protein level was similarly regulated when detected by anti-Polo antibody (Fig. 1F). These results suggest that high basal expression levels and rapid *polo* induction inactivate *lok* in early L3 stage upon irradiation, resulting in the survival of progenitors. Conversely, in late L3 stage, low basal expression and slow *polo* induction activated *lok*, resulting in cell death upon irradiation.

### IR-induced autophagy is necessary for cell death in irradiated late L3 hematopoietic progenitors

We evaluated the level of DNA damage, ROS, and autophagy to determine if they affect the decision between survival and death following IR in early and late L3 progenitors. The extent of DNA damage was determined using an antibody against the phosphorylated form of the histone variant, γ-His2Av, which increases in the vicinity of DSBs. In the absence of IR, the γ-His2Av signal was undetectable in the early and late L3 lymph glands (Fig. S3A). However, 1 h after irradiation, the γ-His2Av signal increased in the MZ and CZ in early and late L3 lymph glands (Fig. S3A). Notably, the intensity of γ-His2Av after irradiation in the early L3 MZ was 4.63-fold higher than that of the late L3 (Fig. S3A). Similarly, when DNA damage was induced by *I-CreI* expression, the intensity of γ-His2Av signal in early L3 progenitors was 1.4-fold higher than that in late L3 progenitors (Fig. S3B). These results suggest that the extent of DNA damage may not explain the survival of early L3 progenitors.

We tested whether ROS affected the decision between the death and survival of progenitors using dihydroethidium (DHE) to detect superoxide radicals. Without irradiation, the amount of ROS in the late L3 progenitors (0.91 ± 0.26) was significantly higher than that in the early L3 progenitors (0.30 ± 0.20) (Fig. 2A upper panels), as previously reported (Owusu-Ansah and Banerjee, 2009). The amount of ROS increased by 1.57-fold in the late L3 progenitors at 1 h after irradiation (1.43 ± 0.44; Fig. 2A). In early L3 progenitors, the DHE intensity increased by 1.9-fold after irradiation; however, the ROS level upon IR was significantly lower than that in the irradiated late L3 progenitors (early L3 vs. late L3; 0.58 ± 0.16 vs. 1.43 ± 0.44, *P<*0.0001, Fig. 2A). These results suggest that ROS may be responsible for the IR-induced death of late L3 progenitors. To test this hypothesis, the antioxidant scavenger protein catalase (*Cat*) was overexpressed in the MZ by *dome-Gal4*. *Cat* expression blocked IR-induced ROS production in the MZ (Fig. S4A) and significantly attenuated IR-induced cell death (Fig. 2B). Moreover, IR-induced cell death was attenuated when the ROS level was decreased by other methods, including overexpression of two extracellular catalases, immune-regulated catalase (*IRC*), and secreted human catalase (*hCatS*), or the RNAi of the transmembrane NADPH oxidase *Duox* (Fig. S4B). These results suggested that IR-induced production of ROS can induce death in the late L3 progenitors.

**Fig. 2.**
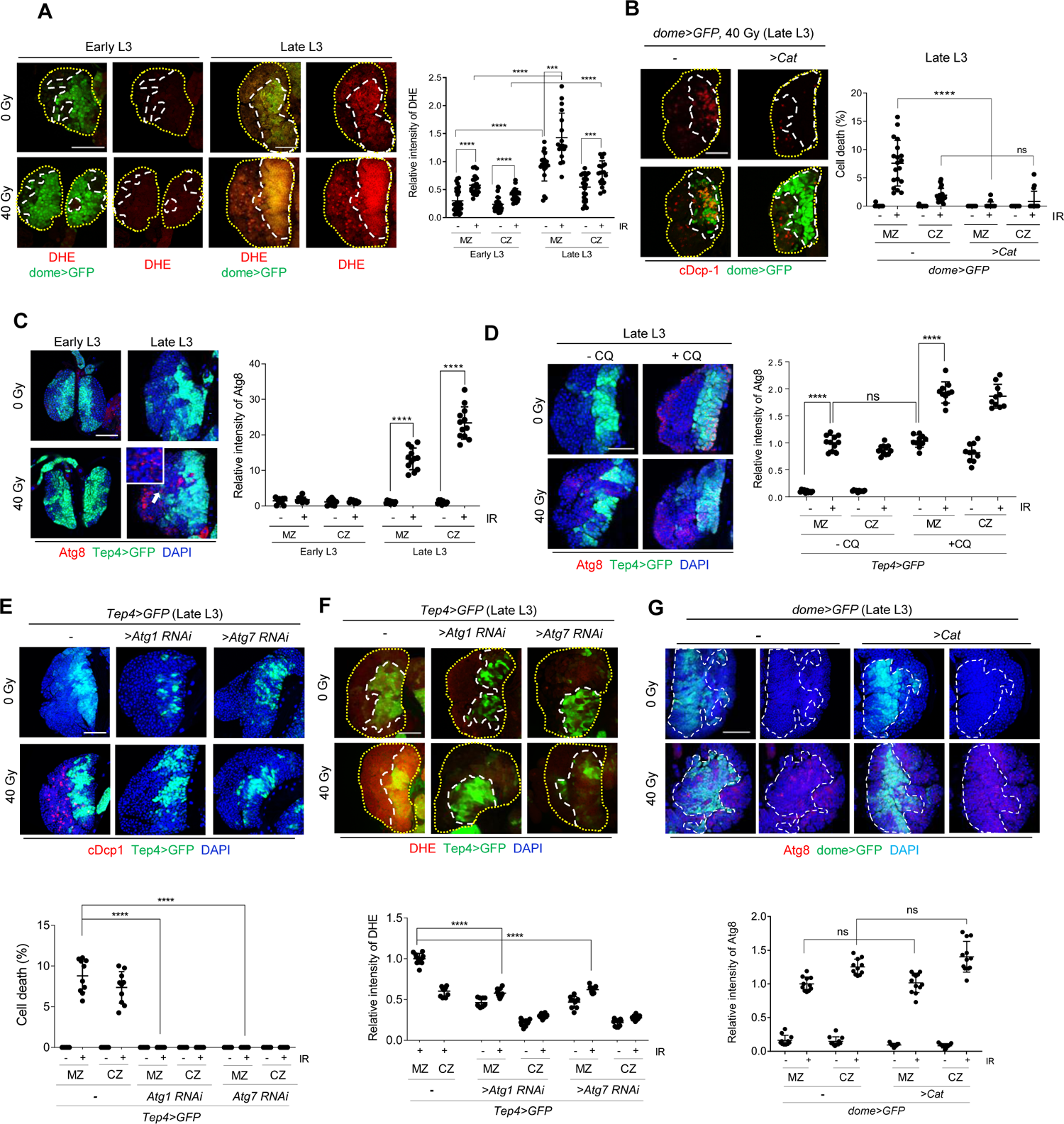
A high level of autophagy and ROS production induced by IR is necessary to induce cell death in the late L3 progenitors. (A) The early and late L3 progenitors were irradiated at 40 Gy. The lymph gland was stained with DHE to detect ROS 1 h after treatment. DHE (red) and dome>GFP (green) indicate ROS and progenitor cells, respectively. Intensities of the DHE signal in progenitors (MZ) and differentiated cells (CZ) with (+) and without (-) IR are shown. The values are normalized relative to the DHE intensity in the late L3 MZ before irradiation in each experiment. Each data point represents a single primary lobe (n>18) and mean±s.d. are shown. **** *P<*0.0001, *** *P<*0.001. (B) The lymph glands from late L3 larvae expressing GFP and Catalase A in MZ (*dome>GFP, CatA*) were stained using cDcp-1 antibody at 4 h after IR. Scale bars, 50 μm. cDcp-1 (red) and dome>GFP (green) indicate apoptotic cells and progenitors, respectively. Percentages of cell number with the cDcp-1 signal in progenitors (MZ) and differentiated cells (CZ) with (+) and without (-) IR are shown. Each data point represents a single primary lobe (n>12) and mean±s.d. are shown. **** *P<*0.0001, ns, not significant. (C) Early and late L3 larvae were irradiated at 40 Gy and 4 h after treatment, and the lymph gland was stained with the Atg8 antibody. DAPI (blue), Atg8 (red), and Tep4>GFP (green) indicate nuclei, autophagosomes, and progenitor cells, respectively. The enlarged image of a cell showing a high number of autophagosomes in irradiated late L3 lymph gland, marked with a white arrow, is shown in the inset. Intensities of the Atg8 signal in progenitors (MZ) and differentiated cells (CZ) with (+) and without (-) IR are shown. The values are normalized relative to the Atg8 intensity in the early L3 MZ before irradiation in each experiment. Each data point represents a single primary lobe (n>12), and mean±s.d. are shown. **** *P<*0.0001. (D) The lymph gland of late L3 larvae treated with CQ was stained with Atg8 antibody to detect autophagosomes at 4 h after 40 Gy irradiation. DAPI (blue), Tep4>GFP (green), and Atg8 (red) indicate DNA, progenitors, and autophagosomes, respectively. Intensities of the Atg8 signal in progenitors (MZ) and differentiated cells (CZ) are shown. Each data point represents a single primary lobe (n=12), and mean±s.d. are shown. **** *P<*0.0001, ns, not significant. (E–G) Late L3 larvae expressing RNAi against Atg1 or Atg7 using Tep4-Gal4 was irradiated with 40 Gy. Lymph glands were stained with cDcp-1 (E) and DHE (F) and the signal was quantitated as described above. (G) The lymph glands from late L3 larvae expressing GFP and Catalase A in MZ (*dome>GFP, CatA*) were stained using Atg8 antibody at 4 h after IR. Scale bars, 50 μm. Atg8 (red) and dome>GFP (green) indicate autophagic cells and progenitors, respectively. Intensities of the Atg8 signal in progenitors (MZ) and differentiated cells (CZ) are shown. Each data point represents a single primary lobe (n>11), and mean±s.d. are shown. **** *P<*0.0001, ns, not significant. The boundaries of the primary lobe and MZ are marked with yellow dotted and white broken lines, respectively. Scale bars, 50 μm.

Next, we evaluated the extent of autophagy after irradiation in the early and late L3 lymph glands using LysoTracker, an acidotropic dye. The intensity of LysoTracker increased in the early and late L3 lymph glands 4 h after irradiation (Fig. S5A), with a higher level in late L3 lymph glands. The induction of autophagy was confirmed by determining autophagosomes through punctate staining of cytoplasmic Atg8 (Yu et al., 2021). Without irradiation, Atg8 staining was undetectable in the early and late L3 lymph glands (Fig. 2C). Additionally, 4 h after irradiation, punctate Atg8 staining increased in the late L3 but not in the early L3 lymph glands (Fig. 2C). To further confirm that the increased number of autophagosomes was because of increased autophagic flux rather than reduced consumption of autophagosomes by lysosomes, Atg8 staining was performed after the larvae were treated with chloroquine (CQ). When lysosomal activity was blocked by CQ, the Atg8 signal was significantly increased (Fig. 2D), confirming that IR increased autophagy in the late L3 lymph gland. When autophagy was blocked by RNAi expression against *Atg1* or *Atg7* using *Tep4-Gal4* (Fig. S5B), IR-induced cell death was not induced in late L3 progenitors (Fig. 2E). Moreover, blocking IR-induced autophagy attenuated ROS production (Fig. 2F); however, blocking IR-induced ROS production by *Cat* expression did not affect autophagy (Fig. 2G). These results suggest that IR-induced autophagy induces the death of late L3 progenitors by increasing ROS levels.

### Enhancing either autophagy or ROS production is sufficient to induce cell death in the late L3 hematopoietic progenitors

To investigate whether enhanced autophagy or ROS production can induce cell death in late L3 progenitors, autophagy or ROS was increased by rapamycin treatment or via RNAi-mediated knockdown of ND75, a component of complex I of the electron transport chain in mitochondria, respectively. Increased autophagy or ROS in late L3 progenitors caused cell death without irradiation (Fig. 3), suggesting that high levels of autophagy or ROS can induce cell death.

**Fig. 3.**
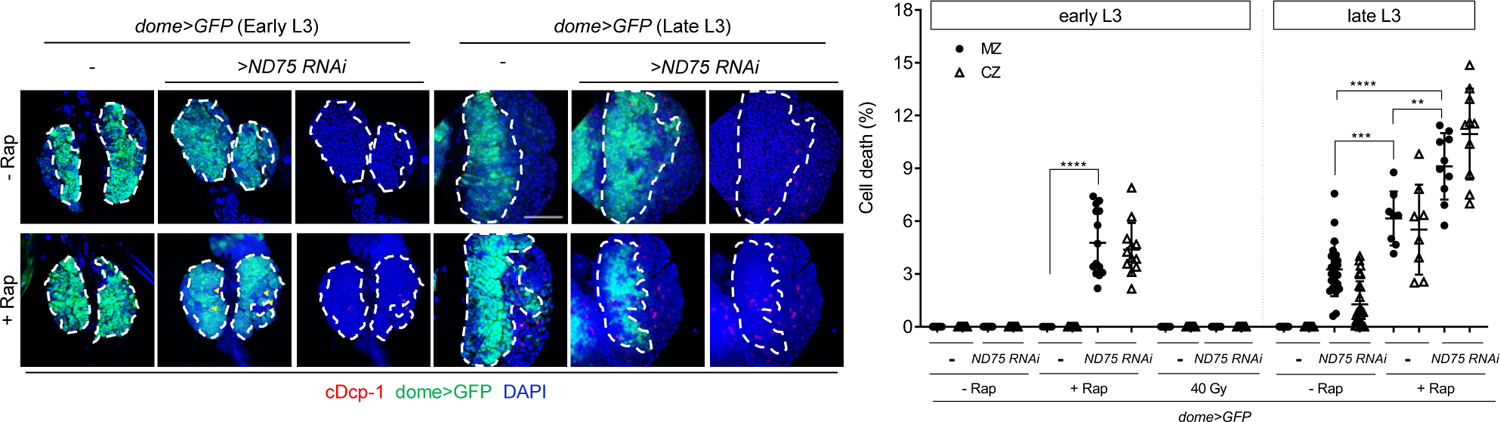
Simultaneous induction of ROS and autophagy is sufficient to induce cell death in early L3 progenitors The early and late L3 larvae expressing ND75 RNAi in MZ (*dome>ND75 RNAi*) were treated with rapamycin (Rap) for 24 h, and the lymph gland was stained with the cDcp-1 antibody. DAPI (blue), dome>GFP (green), and cDcp-1 (red) indicate DNA, progenitors, and apoptotic cells, respectively. Percentages of cell number with cDcp-1 signal in the lymph gland in (A) are shown. Each data point represents a single primary lobe (n>10), and mean±s.d. are shown. **** *P<*0.0001, *** *P<*0.001, ** *P<*0.01.

### Simultaneous increase of ROS and autophagy can induce cell death in the early L3 hematopoietic progenitors

Using *ND75* RNAi or rapamycin treatment, autophagy (Fig. S6) or ROS production (Fig. S4D) in early L3 progenitors was increased to a level comparable to that in irradiated late L3 progenitors, respectively. Unlike late L3 progenitors, individual enhancement of autophagy or ROS was not sufficient to induce cell death in early L3 progenitors (Fig. 3). Conversely, the simultaneous induction of autophagy and ROS significantly increased cell death in early L3 progenitors (Fig. 3). Additionally, the simultaneous induction of autophagy and ROS further increased cell death in the late L3 progenitors compared with individual treatments (Fig. 3). Moreover, rapamycin-induced autophagy was increased by elevating ROS levels via *ND75* knockdown; however, ROS production alone did not affect autophagy in the early and late L3 progenitors (Fig. S6). These results suggest that ROS induced by autophagy in irradiated late L3 progenitors may further increase autophagy, generating a positive feedback loop.

### *polo* can block IR-induced autophagy and ROS production by inactivating *lok*

Atg8 and DHE staining were performed in the early L3 *polo* mutant to determine the genes required for IR-induced autophagy and ROS production. *polo* mutations increased autophagy and ROS production upon IR (Fig. 4A), suggesting that *polo* is required for attenuating IR-induced autophagy and ROS production. IR-induced autophagy and ROS production were significantly attenuated in late L3 *lok^P6^* mutant or when *poloT182D* was overexpressed in the late L3 MZ using *dome-Gal4* (Fig. 4B,D,E). Additionally, *lok* overexpression in early and late L3 progenitors was sufficient to induce autophagy and ROS production (Fig. 4C–E). In contrast, IR-induced autophagy and ROS production in the *dp53^5A-1-4^* mutant were comparable to those in the control (Fig. S7). Furthermore, IR-induced autophagy and ROS production in the early L3 *polo* mutant were significantly attenuated in the *lok polo* double mutant (Fig. 4A), confirming that *polo* can block IR-induced autophagy and ROS production by negatively regulating *lok*.

**Fig. 4.**
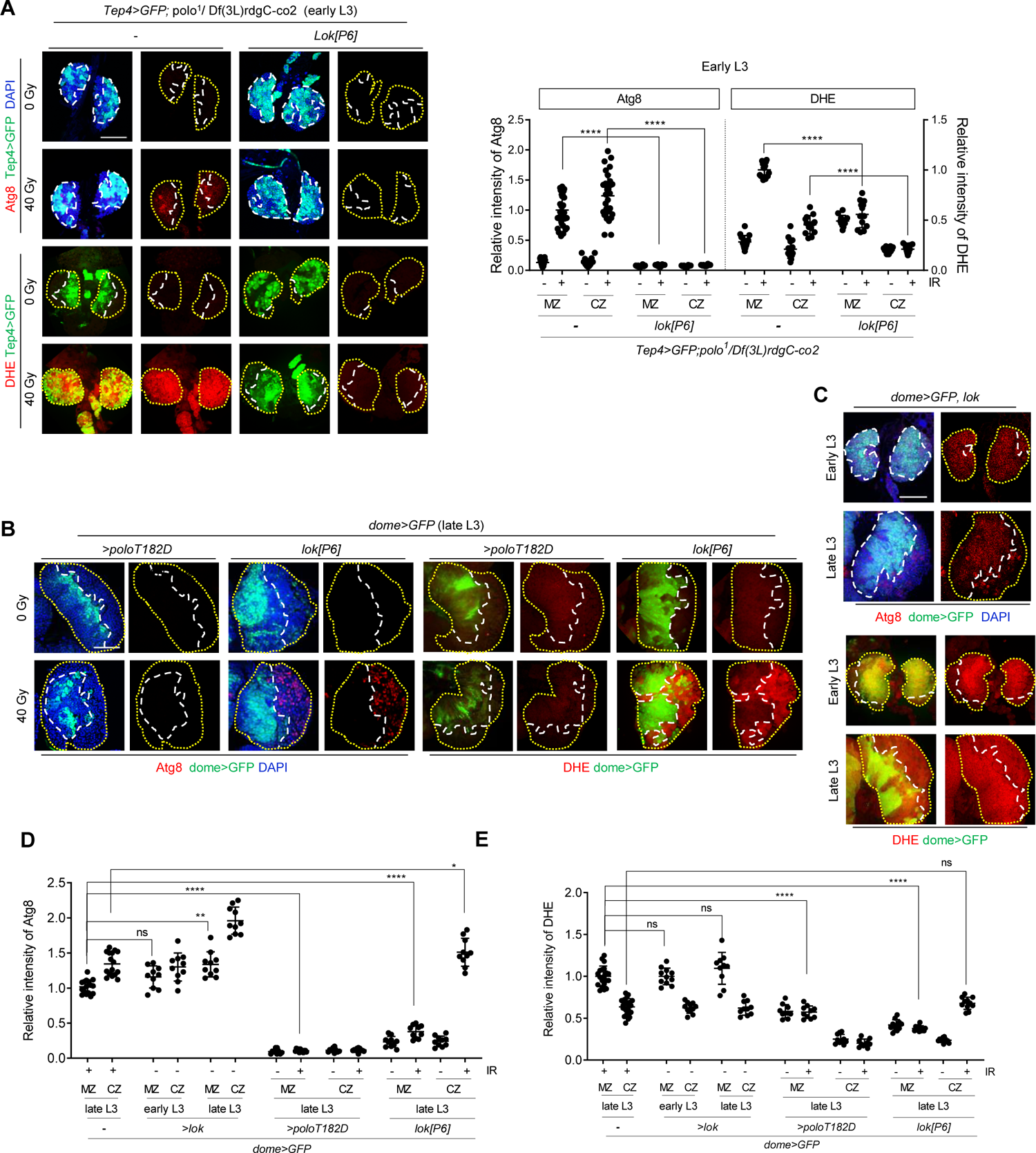
*polo* can block IR-induced autophagy and ROS by inactivating *lok* The larvae with the following genotypes were irradiated at 40 Gy: early L3 *polo* single and *lok polo* double mutant (A), late L3 larvae expressing *poloT182D* in the MZ (B), late L3 *lok* mutant (B), and early and late L3 larvae expressing *lok* in the MZ (C). The lymph glands were stained with the Atg8 antibody or DHE at 4 h or 1 h after 40 Gy irradiation, respectively. DAPI (blue), Tep4>GFP (green), Atg8 (red), and DHE (red) indicate DNA, progenitors, autophagosomes, and ROS, respectively. Scale bars, 50 μm. (A) The relative intensities of Atg8 and DHE in samples (A) were quantified. Each data point represents a single primary lobe (n>10) and mean±s.d. are shown. **** *P*<0.0001. (D, E) The relative intensities of Atg8 (D) and DHE (E) in samples (B, C) were quantified. Each data point represents a single primary lobe (n>10) and mean±s.d. are shown. **** *P<*0.0001, ** *P<*0.01, * *P<*0.05, ns, not significant.

### Hematopoietic progenitors irradiated at early L3 stage undergo cell cycle arrest and premature differentiation

Since the hematopoietic progenitors irradiated during early L3 stage did not exhibit cell death, their fate was investigated further. To determine the short-term effect of irradiation, cell cycle arrest was assessed by staining mitotic cells with a phospho-histone H3 (PH3) antibody 1 h after IR. The percentage of mitotic cells in the MZ was significantly reduced upon irradiation (Fig. 5) (from 11.71 ± 5.02% to 0.80 ± 1.25%). This result suggested that hematopoietic cells in the early L3 lymph gland were arrested at the G2 phase after high-dose irradiation, assuming that the length of the G2 phase of these cells was less than 1 h.

**Fig. 5.**
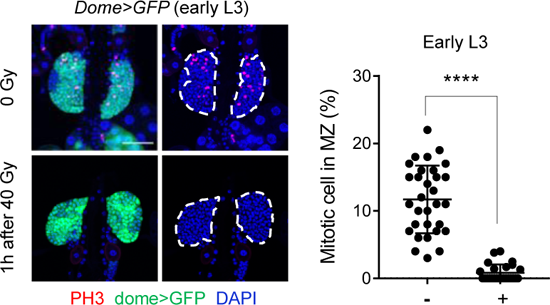
IR induces cell cycle arrest of progenitors in the early L3 lymph gland The early L3 larvae were irradiated at 40 Gy, and the lymph gland was stained using phospho-histone H3 (PH3) antibody to mark mitotic cells at 1 h post-irradiation. DAPI (blue), dome>GFP (green), and PH3 (red) indicate DNA, progenitors, and mitotic cells, respectively. The boundary of the dome>GFP-stained MZ is marked with white broken lines. Scale bars, 50 μm. The percentages of the number of PH3-positive cells in MZ in early L3 lymph gland with (40 Gy) and without (0 Gy) IR are shown. Each data point represents a single primary lobe (n>28), and mean±s.d. are shown. **** *P<*0.0001.

To understand the long-term effect of irradiation on surviving hematopoietic progenitors, differentiation was determined at 40 h post-irradiation by staining progenitors, lamellocytes, crystal cells, and early-stage plasmatocytes with *dome>GFP*, phalloidin, Hindsight (Hnt), and peroxidasin (Pxn), respectively (Fig. 6). In wild type lymph gland, although the size of the primary lobe was not changed by IR (Fig. S8A), irradiation decreased the area of *dome>GFP*-positive progenitors (Fig. S8B) and the intensity of *dome>GFP* (Fig. S8C) after 40 h. Lamellocytes, undetected under normal conditions, were detected post-irradiation in 79.7 ± 11% of the lymph gland (Fig. 6A,B). Furthermore, the percentages of crystal cells (2.2 ± 0.7% at 0 Gy and 3.5 ± 0.6% at 40 Gy) and plasmatocytes (44.2 ± 8.7% at 0 Gy and 75.0 ± 8.6% at 40 Gy) were increased by 1.6- and 1.7-fold, respectively (Fig. 6A,C,D). These results demonstrated that irradiation in early L3 larvae induced premature differentiation of hematopoietic progenitors into three hemocyte cell types.

**Fig. 6.**
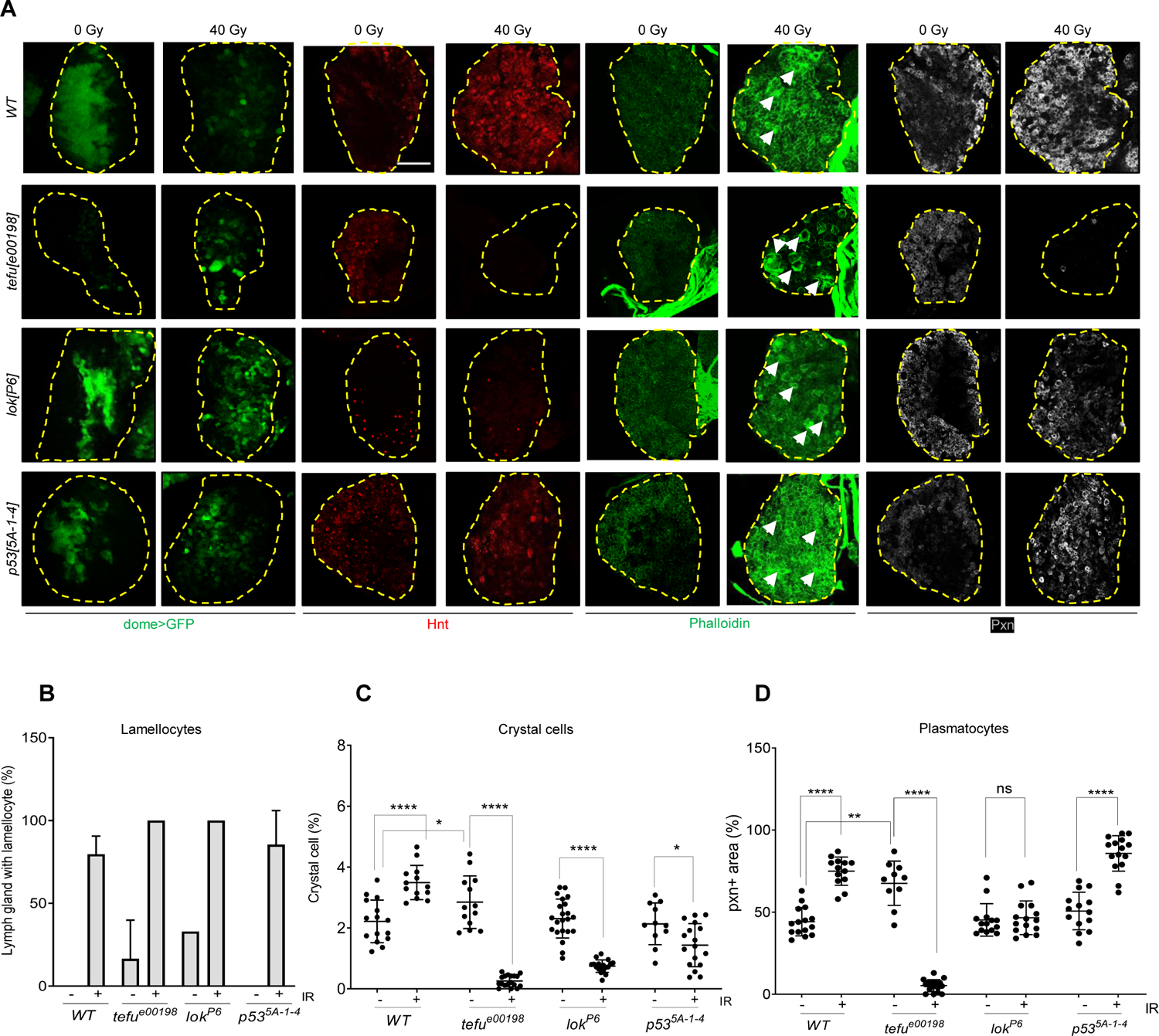
IR induces premature differentiation of progenitors when irradiated in early L3 larvae. (A) The early L3 larvae of wild type, *tefu^e00198^*, *lok^P6^*, and *dp53^5A-1-4^* mutants were irradiated at 40 Gy, and the lymph gland was stained with dome>GFP, hindsight (Hnt), phalloidin, and peroxidasin (Pxn) 40 h after treatment. The boundary of the primary lobe is marked with yellow broken lines. Scale bars, 50 μm. Dome>GFP (green), Hnt (red), phalloidin (green), and Pxn (white) indicate progenitors, crystal cells, F-actin-rich lamellocytes, and early stage of plasmatocytes, respectively. Arrows indicate lamellocytes. The representative images of the maximum projection of entire z-stacks of lymph glands are shown for phalloidin and Hnt. Single middle z-stack images are shown for dome>GFP and Pxn. (B–D) The quantitation of hemocytes in (A). The percentages of primary lobes containing lamellocytes (B), the number of crystal cells (C), and the Pxn-positive area (D) with (+) and without (-) IR are shown. Each data point represents a single primary lobe (n>10), and mean±s.d. are shown. **** *P<*0.0001, ** *P<*0.01, * *P<*0.05 ns, not significant.

To determine whether premature differentiation by IR was induced by DNA damage, the differentiation of hemocytes was assessed after generating DSBs by heat-shock-driven expression of *I-CreI*. Similar to irradiation, *I-CreI* expression significantly induced lamellocyte differentiation (85.0 ± 26.0%) (Fig. S9A,B). In contrast, the proportion of crystal cells decreased by 54.2% after I-CreI expression (from 16.8 ± 4.5% without heat shock to 9.1 ± 3.7% after heat shock) (Fig. S9A,C). Additionally, large Pxn-positive plasmatocyte areas were detected in the absence of heat shock (68.5 ± 11.9%), which persisted after *I-CreI* expression (66.1 ± 9.1%) (Fig. S9A,D). As a control, heat-shock treatment of wild type L3 larvae did not affect differentiation (Fig. S9). These results suggested that DNA damage may induce the differentiation of lamellocytes but not of crystal cells and plasmatocytes.

### Genes involved in IR-induced premature differentiation of progenitors in early L3 stage

The roles of *tefu, dp53,* and *lok* were investigated to determine the genes required for the IR-induced premature differentiation of hematopoietic progenitors in the early L3 lymph gland. As shown in our previous study, without irradiation, the size of the lymph gland in *tefu^e00198^* was half that of the control (Fig. S8A) (Nguyen et al., 2021). The size of *lok^P6^* and *dp53^5A-1-4^* was similar to that of the wild type without irradiation in the present study (Fig. S8A). Upon irradiation, the size of the lymph glands was unchanged in the wild type and all three mutants (Fig. S8A). In response to irradiation, the number of lymph glands containing F-actin-rich lamellocytes was increased in *tefu^00198^, lok^P6^*, and *dp53^5A-1-4^* mutants, similar to those in the wild type (Fig. 6A,B), suggesting that *tefu, lok*, and *dp53* are not required for IR-induced differentiation of lamellocytes. Contrary to the wild type, which showed a 1.57-fold increase in crystal cell differentiation after 40 Gy irradiation (from 2.22 ± 0.70% without IR to 3.49 ± 0.56% after IR), the percentage of crystal cells after IR in *tefu^00198^, lok^P6^,* and *dp53^5A-1-4^* mutants was reduced to 10.7%, 30.4%, and 66.7% of the unirradiated sample, respectively (from 2.8 ± 0.9%, 2.3 ± 0.6%, and 2.1 ± 0.7% without IR to 0.3 ± 0.2%, 0.7 ± 0.2%, and 1.4 ± 0.7% after IR in *tefu^00198^*, *lok^P6^*, and *dp53^5A-1-4^* mutants, respectively) (Fig. 6A,C). This result suggests that *tefu* and *lok* may play a major role in inhibiting the differentiation of crystal cells upon irradiation. Regarding IR-induced differentiation into plasmatocytes, a 1.71- and 1.69-fold increase in the Pxn-positive area was observed after irradiation in the wild type and *dp53^5A-1-4^* mutant, respectively, suggesting that *dp53* is not required for this process (wild type: 44.2 ± 8.67% at 0 Gy and 75 ± 8.6% at 40 Gy; *dp53^5A-1-4^*: 50.7 ± 11.4% at 0 Gy and 85.8 ± 10.8% at 40 Gy) (Fig. 6A,D). In contrast, the percentage of plasmatocytes was unchanged after irradiation in *lok^P6^* (45.3 ± 9.9% at 0 Gy and 46.6 ± 10.3% at 40 Gy) (Fig. 6A,D), suggesting that *lok* is required for IR-induced premature differentiation of plasmatocytes. In *tefu^e00198^*, the Pxn-positive area at 0 Gy (67.6 ± 13.5%) was severely decreased (5.3 ± 3.7%) at 40 Gy (Fig. 6A,D), suggesting that *tefu* may play a role in inhibiting the differentiation of plasmatocytes in response to IR. Notably, the percentage of progenitors and the intensity of *dome>GFP* were severely lower in *tefu^e00198^* without irradiation than in the wild type (Fig. S8B,C). Moreover, they increased in *tefu^e00198^* after 40 Gy, in contrast to those in the wild type.

## Discussion

We investigated the cell fate of *Drosophila* hematopoietic progenitors during larval development and determined the mechanism underlying the decision between death and survival after IR. Upon IR, the cell fate of larval hematopoietic progenitors changed from survival in early L3 stage to death in mid/late L3 stage. Furthermore, the expression level of *polo* and the high basal levels of ROS in late L3 progenitors regulated the decision between survival and death. A low basal level of *polo* in late L3 stage enabled IR-induced *lok* activation, which induced autophagy and increased ROS production, resulting in cell death. In contrast, a high basal level of *polo* expression in early L3 stage blocked IR-induced *lok* activation, enabling the survival of irradiated progenitors. The surviving progenitors underwent transient cell cycle arrest and premature differentiation regulated by *tefu* and *lok*.

The percentage of *polo*-expressing cells and *polo* expression level in early L3 progenitors were higher than those in late L3 progenitors. Notably, during *Drosophila* larval hematopoiesis, progenitor cells proliferate in the early larval stages (51% in S, 19% in G2, and 8% in M) and become quiescent in late L3 stage (38% in S, 57% in G2, and 0% in M) where they are arrested at the G2 phase (Krzemien et al., 2010; Sharma et al., 2019). As a key cell cycle regulator, PLK1 expression is regulated during the cell cycle, such that it begins to increase from the S/G2 phase and peaks at the M phase (Golsteyn et al., 1995; Golsteyn et al., 1994), which suggests that changes in *polo* level during development may reflect the cell cycle profile. In response to IR, *polo* transcription appeared to increase via a separate mechanism because its expression was further increased 1 h after IR at which point the percentage of mitotic cells was significantly decreased (Figs. 2D, 5). Moreover, the *cis*-elements required for regulating *polo* expression during larval development and in response to IR were present in the genomic sequence of GFP-polo (Fig. 2D–G), encompassing approximately 7 kb, including the *polo* gene and 5′ and 3′ UTR (Moutinho-Santos et al., 1999). Additionally, several putative binding sites for Ftz, Zeste, Prd, DSXF, DSXM, Hb, Mad, B factor, Dfd, and TII were identified in this reqion via *in silico* analysis. However, their roles in developmental and IR-induced regulation of *polo* transcription remains undetermined.

During DDR, PLK1 activity is regulated to perform multiple functions. When DNA damage occurs in interphase, PLK1 is inhibited in an AMT/ATR-dependent manner to prevent entry into mitosis (Smits et al., 2000; Tsvetkov and Stern, 2005). When DNA damage is repaired, PLK1 activation is required for checkpoint recovery to enable cells to resume the cell cycle (van Vugt et al., 2004). Notably, the deregulation of PLK1 activity results in tumorigenesis, and PLK1 is overexpressed in various cancers (Iliaki et al., 2021). Moreover, PLK1 overexpression is associated with poor prognosis and resistance to chemotherapy. Consistent with our results, PLK1 inhibition in breast cancer cells reduced radiation-induced ROS production and autophagy, resulting in enhanced radiosensitivity (Wang et al., 2021). However, the role of PLK1 during autophagy induction and apoptosis remains controversial as it shows pro-autophagic (Ruf et al., 2017), anti-autophagic (Tao et al., 2017), pro-apoptotic (Jang et al., 2011), and anti-apoptotic effects (Matthess et al., 2014), possibly owing to different cell types or specific experimental conditions. Thus, our study on larval hematopoietic progenitors provided an *in vivo* model to confirm that *polo* played anti-autophagic and anti-apoptotic roles.

*polo* enabled the survival of irradiated early L3 hematopoietic progenitors by negatively regulating *lok*. Although direct interaction between *lok* and *polo* has not been investigated in *Drosophila*, PLK1 directly binds, phosphorylates, and inactivates CHK2 in mammals (Tsvetkov et al., 2005; van Vugt et al., 2010). Moreover, the interaction between PLK1 and CHK2 is reportedly cell cycle-dependent; they dissociate in response to DNA damage in the G2 and M phases but not in the S phase (Tsvetkov et al., 2005). Together with the high level of *polo* in early L3 progenitors, the active proliferation status of early L3 progenitors may further inactivate *lok*. Besides its role in phosphorylating and activating *dp53*, which results in the induction of pro-apoptotic genes (*reaper* and *hid*), *lok* was necessary and sufficient to induce autophagy and ROS production in L3 hematopoietic progenitors. However, autophagy and ROS production by *lok* was not mediated by *dp53*, since IR-induced ROS production and autophagy occurred in *dp53* mutants (Fig. S7). Besides the ATM-mediated post-translational modification pathway during acute autophagy induction in response to DNA damage (Chen et al., 2015), the role of CHK2 in autophagy induction has recently been reported. Under various stress conditions, CHK2 phosphorylates Beclin 1 (Guo et al., 2020), FOXK (Chen et al., 2020), and ULK1 (mammalian Atg1) (Guo et al., 2022) and induces autophagy. Nevertheless, whether Atg6 (*Drosophila* Beclin 1), FoxK (*Drosophila* FOXK), and Atg1 (*Drosophila* ULK1) are physiological substrates of *lok* to regulate autophagy remains to be studied.

We detected four different ROS levels during development of *Drosophila* hematopoietic progenitors in response to IR. The lowest ROS level was detected in unirradiated early L3 progenitors, which slightly increased upon irradiation. Furthermore, in the late L3 progenitors, the basal level of ROS was higher than that in irradiated early L3 progenitors and functioned as signaling molecules to induce differentiation. Upon IR, ROS levels in late L3 progenitors further increased, resulting in cell death. The induction of autophagy alone was sufficient to induce cell death in late L3 progenitors exhibiting the highest ROS level, but not in the irradiated early L3 progenitors with lower ROS levels (Figs. 2A, 3). This result suggests that a threshold ROS amount and autophagy induction is required to induce cell death and that the amount of ROS in irradiated early L3 progenitors was not sufficient. Therefore, the simultaneous induction of autophagy and ROS production is required to induce cell death in early L3 progenitors. In contrast, the high basal level of ROS in late L3 progenitors was sufficient to induce cell death when only autophagy was induced. This result suggests that the basal level of ROS is likely a critical factor affecting the role of autophagy during radiation therapy.

In late L3 progenitors, autophagy was required to promote IR-induced death and was sufficient to induce cell death when the ROS level was high, suggesting the cytotoxic role of autophagy. Radiation-induced autophagy is generally considered cytoprotective, such that inhibition of autophagy sensitizes tumor cells to death upon radiation therapy (Gewirtz, 2014). However, previous studies suggest that autophagy has a cytotoxic effect, especially when irradiated cells are treated with radiosensitizing agents (Cao et al., 2006; Fujiwara et al., 2007; Gewirtz, 2014). In late L3 progenitors, autophagy induction by rapamycin treatment resulted in cell death, indicating that it can be defined as autophagy-mediated cell death according the current guidelines of scientific communities (Denton and Kumar, 2019). Additionally, we found that IR-induced ROS production in L3 progenitors was attenuated by inhibiting autophagy (Fig. 2F), suggesting that autophagy acts upstream of ROS during IR-induced cell death. Consistent with our finding, the caspase inhibitor, z-VAD, mediated cell death in mouse L929 cells only when autophagy occurred, as ROS is increased to cytotoxic levels via selective autophagic degradation of catalase (Yu et al., 2006). Although the regulation of autophagy by oxidative stress is well established (Redza-Dutordoir and Averill-Bates, 2021), IR-induced autophagy in L3 progenitors was not affected by inhibited ROS production (Fig. 2G). Instead, when ROS was induced in the presence of rapamycin-induced autophagy, autophagy was further increased (Fig. S6), thus generating a positive feedback regulation.

Surviving progenitors irradiated at early L3 stage underwent cell cycle arrest (Fig. 5) followed by premature differentiation (Fig. 6). In response to environmental stresses, including hypoxia, starvation, infection, and oxidative stress, *Drosophila* larval lymph glands fail to maintain homeostasis, resulting in premature differentiation and depletion of progenitor populations (Sinenko et al., 2011). IR-induced differentiation of crystal cells and plasmatocytes were affected by *tefu* and *lok* mutation. Severe reduction of crystal cells and plasmatocytes in irradiated *tefu* mutants suggest that *tefu* may be necessary for the survival or proliferation of crystal cell/plasmatocyte precursors. In the *Drosophila* germline cells, *lok* negatively regulates *bam* protein expression (Ma et al., 2016). *microRNA-7* interacts with *bam* to maintain the level of hematopoietic progenitor cells in the *Drosophila* lymph gland via negative regulation of *yan*, which generally promotes blood cell differentiation (Tokusumi et al., 2011). Therefore, *lok* activity in the irradiated early L3 progenitors may decrease *bam* levels, thereby increasing *yan* activity and promoting crystal cell and plasmatocyte differentiation. In mammals, lymphoid differentiation of hematopoietic stem cells (HSCs) are induced by BATF in a G-CSF/STAT3-dependent manner (Wang et al., 2012). Precocious differentiation of HSCs after DNA damage occurs rapidly post-injury and may account for rapid blood lineage regeneration in an attempt to prevent the accumulation of HSCs with genomic abnormalities and to return the system to homeostasis (Wang et al., 2012).

Overall, the *polo* expression level was critical for cells to decide between cell survival and death upon irradiation via the negative regulation of *lok* activity that induces autophagy and ROS production. Furthermore, in this biological context, autophagy induction and high ROS production levels were necessary for inducing cell death. Although the mechanisms underlying the regulation of *polo* expression, *lok*-mediated autophagy and ROS production, and autophagy-mediated cell death remains elusive, we provide a model system to study the cell fate decision of stem cells upon IR during development (Fig. 7). Since PLK1 (Iliaki et al., 2021), ROS (Pelicano et al., 2004), and autophagy levels (Li et al., 2020) are altered in cancer cells, the *Drosophila* lymph gland will be a valuable *in vivo* model to investigate the functional mechanism of *polo*, *lok*, ROS, and autophagy during cell death and provide clues for developing efficient anti-cancer drugs.

**Fig. 7.**
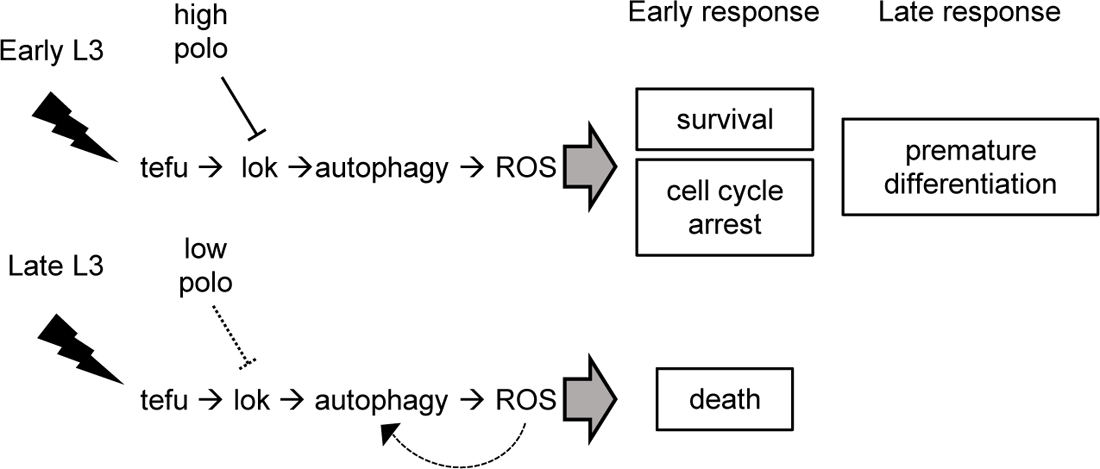
Proposed mechanism of how IR-induced cell survival and death is regulated in early and late L3 progenitors. In the late L3 progenitors, IR-induced activation of *lok* induces autophagy and ROS production, resulting in cell death. In the early L3 progenitors, a high level of *polo* blocks IR-induced lok activation, resulting in cell survival, cell cycle arrest, and premature differentiation.

## Materials and Methods

### Drosophila melanogaster

All *Drosophila* fly stocks were reared on cornmeal, agar, dextrose, and yeast medium under a 12 h light: 12 h dark cycle at 50% relative humidity and 25 °C in an incubator unless otherwise specified. Canton S was used as the wild type. Genotypes for each figure and number of samples investigated are listed in Table S1. The following *Drosophila* stocks were used in this study: *p53^5A-1-4^* (BL6815), *hs-I-CreI* (BL6937), *UAS-Cat* (BL24621), *UAS-poloT182D* (BL8434), *polo RNAi* (BL33042), *GFP-polo* (BL84275), and *GMR-gal4/CyO* (BL9146) were obtained from the Bloomington Drosophila Stock Center (Bloomington, IN, USA). *ND75* RNAi (NIG2286R-3) was obtained from the National Institute of Genetics (Japan). *Tep4-Gal4* (DGRC105442), *tefu^GS13617^* (DGRC204829), *polo^1^* (DGRC105973), and *Df(3L)rdgC-co2* (DGRC106730) were obtained from the Kyoto Stock Center (Japan). Other fly stocks were *lok^P6^* (W. Theurkauf); *dome-Gal4* (M. Zeidler); *Hml-Gal4* (S. Sinenko); *dome-MESOGal4, UAS-EYFP*, and *Antp-gal4* (U. Banerjee); *UAS-lok* and *UAS-lok-KD* (Y.-H. Song); *UAS-IRC, UAS-hCatS,* and *UAS-Duox RNAi #44* (W.J. Lee); and *tefu^e00198^* (Exelixis, Boston, MA, USA).

### Rapamycin treatment

Rapamycin (LC Laboratories, Woburn, MA, USA) was dissolved in ethanol and mixed with the media when preparing food dishes. The final rapamycin dose was 2 µM, and equal volumes of ethanol alone were added to the vehicle control food. Next, the larvae were reared on normal food and transferred to rapamycin food plates for 24 h before dissection.

### Chloroquine treatment (autophagic flux assay)

Chloroquine diphosphate salt (CQ, Sigma-Aldrich, St. Louis, MO, USA) was dissolved in distilled water and mixed with the media when preparing food dishes. The final CQ dose used was 10 mg/mL. Next, the larvae were reared on normal food and transferred to CQ food plates for 12 h before dissection.

### Immunofluorescence staining and antibodies

The immunofluorescence staining protocol used was previously described by Nguyen et al. (2021). The following primary antibodies were used in this study: rabbit anti-cleaved Dcp-1 (Asp215) (1:250, Cell Signaling, Danvers, MA, USA), rabbit anti-peroxidasin (Pxn, 1:2500) (Yoon et al., 2017), mouse anti-Hindsight (Hnt, 1: 20, Developmental Studies Hybridoma Bank, Iowa City, IA, USA), Phalloidin-FITC (1:200, Sigma P5282), rabbit anti-phospho-Histone H3 (Ser10) (1:300, Merck, Damstadt, Germany), rabbit anti-GABARAP+GABARAPL1+GABARAPL2 (Atg8, 1:400, Abcam, Cambridge, UK), rabbit anti-histone H2AvD pS137 (γ-His2-Av, 1: 400, Rockland Immunochemicals, Pottstown, PA, USA), rabbit anti-GFP (1:200, Abcam), mouse anti-GFP (1:100, Santa Cruz Biotechnology, Dallas, TX, USA), rabbit anti-Polo (1:100) (Arnaoutov et al., 2020), goat anti-rabbit Alexa Fluor 568, goat anti-rabbit Alexa Fluor 647, goat anti-mouse Alexa Fluor 488, goat anti-rabbit Alexa Fluor 488, and goat anti-mouse Alexa Fluor 568 (Invitrogen, Waltham, MA, USA).

### Lysotracker staining

The lymph glands were dissected at 4 hours after IR. Lysotracker staining was performed using LysoTracker Red DND-99 (Molecular Probes, Inc., Eugene, OR, USA, L7528), as previously described (DeVorkin and Gorski, 2014). First, the dissected lymph glands were stained with 10 µM LysoTracker Red DND-99 solution in phosphate-buffered saline (PBS), incubated in the dark for 3 min at room temperature, and washed in PBS. Next, the samples were fixed in 4% formaldehyde for 20 min and washed with 0.4% and 0.1% PBT. Afterward, the samples were stained with 4′,6-diamidino-2-phenylindole (DAPI) and mounted. All samples were immediately visualized.

### ROS staining

The lymph glands were dissected at 1 hours after IR. ROS staining was performed using DHE (Molecular Probe #D11347) as previously described (Evans et al., 2014). First, the dissected lymph glands were stained with 30 µM DHE solution in Schneider’s medium, incubated in the dark for 5 min at room temperature, and washed in Schneider’s medium. Next, the samples were fixed in 7% methanol-free formaldehyde for 5 min and washed with PBS. Afterward, the samples were stained with DAPI and mounted. All samples were immediately visualized using a confocal laser scanning microscope (LSM 700, Carl Zeiss, Oberkochen, Germany).

### RNA FISH and protein immunofluorescence (IF) double labeling

To generate RNA probes, a portion of the *polo* cDNA (291 bp; forward primer: GCA ATC TCT TCC TCA ACG, reverse primer: GTT TCC TTA AGT AGC TGG GC) was cloned into the pGEM-T easy vector (Promega, Madison, WI, USA) in sense and antisense orientations. Next, the plasmid DNA was linearized by SalI digestion and transcribed by T7 RNA polymerase using Digoxigenin (Dig) RNA Labeling Mix (Roche Diagnostics, Germany) for 2 h at 37 °C. The reaction was stopped by adding 0.2 M EDTA (pH 8.0), and the RNA was precipitated by ethanol and dissolved in RNase-free distilled water. Subsequently, FISH/IF double labeling was performed to detect the *polo* transcript and GFP as previously described with slight modifications (Lécuyer et al., 2008; Park et al., 2019). Briefly, early and late L3 larvae were irradiated at 40 Gy, and lymph glands were dissected in 1× PBS after 1, 2, and 4 h. Thereafter, the samples were fixed in a 1:1 mixture of 4% para-formaldehyde (Sigma-Aldrich) and heptane (Sigma-Aldrich) for 20 min with shaking. Next, the samples were permeabilized with 3 μg/mL proteinase K for 13 min at room temperature and incubated on ice for 1 h before washing with 2 mg/mL glycine solution. After re-fixation and blocking, the samples were hybridized with a 1 μg/mL Dig-labeled RNA probe at 56 °C overnight. The lymph glands were incubated with biotin-conjugated mouse monoclonal anti-DIG (1:400, Jackson ImmunoResearch Lab Inc., West Grove, PA, USA) and rabbit anti-GFP antibodies (1:200, Abcam) for 2 h at room temperature with shaking. Afterward, secondary antibody incubation was performed for 1 h with 1× streptavidin-HRP (Invitrogen) and goat anti-rabbit Alexa Fluor 488 (1:200, Invitrogen) at room temperature. Moreover, signal amplification was performed with Alexa Fluor 555 Tyramide conjugate (1:50) in 1× reaction buffer supplied with Alexa Flour 555 Tyramide SuperBoost kit (Invitrogen) for 1 h at room temperature. The reaction was terminated by adding a stop solution for 10 min, and the lymph glands were mounted in 0.5% n-propyl gallate dissolved in glycerol.

### Quantification and statistical analysis

#### Quantification of cell death

TepIV-Gal4- or dome-Gal4-driven GFP was used as a marker for MZ. The total number or size of the lobes was measured using DAPI staining. Images were processed using ImageJ (NIH, Bethesda, MD, USA), and the total number of cells in each compartment or total area was determined as previously described (Nguyen et al., 2021).

#### Quantification of the relative intensity of the γ-His2Av signal

GFP driven by dome-Gal4 was used as a marker for MZ. The Zen software (blue edition) defined the regions of interest (ROIs) corresponding to each DAPI-stained cell. One middle confocal section was analyzed in each primary lobe. The average mean intensity of γ-His2Av signal in 20 DAPI-stained cells was quantified in each lobe. Values are shown relative to the γ-His2Av intensity in the late L3 MZ after irradiation in each experiment. At least 17 primary lobes from three independent experiments were analyzed for each sample.

#### Quantitation of the relative intensity of the DHE and Atg8 signal

GFP driven by dome-Gal4 (or Tep4-Gal4) and Hml-Gal4 was used as a marker for MZ and CZ, respectively. The Zen software (blue edition) was used to define the ROIs corresponding to the entire primary lobe and the MZ and CZ areas. One middle confocal section was analyzed in each primary lobe. The average mean intensity of the DHE or Atg8 signal in the 20 cells was quantified for each lobe. The values are shown relative to the DHE or Atg8 intensity in the late L3 MZ before irradiation for each experiment. For the DHE signal, at least 18 primary lobes from three independent experiments were analyzed for each sample. For the Atg8 signal, at least 10 primary lobes from two independent experiments were analyzed for each sample.

#### Quantitation of the relative intensity of GFP-polo, Polo, and polo transcripts

The Pxn and GFP driven by Tep4-Gal4 were used as a marker for CZ and MZ, respectively. The Zen software (blue edition) was used to define the ROIs corresponding to the entire primary lobe and the CZ area. One middle confocal section was analyzed in each primary lobe. The average mean intensity of the Polo, GFP-polo, or *polo* transcript signal in 20 cells was quantified for each lobe. The values are shown relative to the Polo, GFP-polo, or *polo* transcript intensity in the late L3 MZ before irradiation for each experiment. At least 10 primary lobes from two independent experiments were analyzed for each sample.

#### Quantitation of mitotic cells

The percentage of mitotic cells in the lymph gland was calculated by dividing the number of PH3-positive cells by the total number of DAPI-stained cells in the lymph gland. The Fiji software was used to define the ROIs corresponding to the entire primary lobe. At least 44 primary lobes from three independent experiments were analyzed for each sample.

#### Quantitation of crystal cells

The crystal cells were marked by Hnt antibody. The percentage of crystal cells corresponds to the number of crystal cells in the entire section of each primary lobe divided by the total area of the primary lobe. The entire section of each primary lobe was examined using a fluorescent microscope. At least 10 primary lobes from two independent experiments were analyzed for each genotype.

#### Quantitation of plasmatocytes

Early-stage plasmatocytes were marked by the Pxn antibody. The percentage of plasmatocyte area corresponds to the Pxn-positive area divided by the total area of the primary lobe. At least 10 primary lobes from two independent experiments were analyzed for each genotype.

#### Quantitation of lamellocytes

Phalloidin-FITC was used to mark lamellocytes, which are large F-actin-rich flat cells. The percentage of lymph glands containing lamellocytes over the total number of lymph glands was calculated. At least 10 primary lobes from two independent experiments were analyzed for each genotype.

#### Quantitation of progenitors

Progenitors were marked by dome-GFP. The percentage of progenitor area corresponds to the area of the dome-GFP-positive area divided by the total area of the primary lobe.

#### Imaging

Immuno-stained images were visualized using a confocal laser scanning (LSM 700, Carl Zeiss) or fluorescence microscope (IX71, Olympus, Tokyo, Japan). Images were processed using the ImageJ (NIH) and Zeiss Zen software.

#### Statistical analysis

Genotypes for each figure and number of samples tested are listed in Table S1. At least 10 primary lobes from two independent experiments were analyzed for each genotype. The statistical details for each experiment are described in the figure legends. *P*-values are shown in the figures and listed in the figure legends. All statistical analyses were performed using the GraphPad Prism 7 software. The significance of the differences between the two experimental samples was determined using an unpaired t-test with Welch’s correction. Differences were considered statistically significant at *P<*0.05.

## Acknowledgments

This research was supported by the Basic Science Research Program through the National Research Foundation of Korea (NRF) funded by the Ministry of Education (NRF-2020R1I1A3057944).

## Author contributions

Y.-H.S conceived and supervised experiments, wrote the manuscript, and secured funding. T.T.N.T. performed experiments and analyzed data. R.W. performed experiments. G.-C.C. and J.S provided reagents, expertise, and feedback.

## References

Arnaoutov, A., Lee, H., Plevock Haase, K., Aksenova, V., Jarnik, M., Oliver, B., Serpe, M. and Dasso, M. (2020). IRBIT Directs Differentiation of Intestinal Stem Cell Progeny to Maintain Tissue Homeostasis. iScience 23, 100954.

Azad, M. B., Chen, Y. and Gibson, S. B. (2009). Regulation of autophagy by reactive oxygen species (ROS): implications for cancer progression and treatment. Antioxid Redox Signal 11, 777–790.

Banerjee, U., Girard, J. R., Goins, L. M. and Spratford, C. M. (2019). Drosophila as a Genetic Model for Hematopoiesis. Genetics 211, 367–417.

Barazzuol, L., Ju, L. and Jeggo, P. A. (2017). A coordinated DNA damage response promotes adult quiescent neural stem cell activation. PLoS Biol 15, e2001264.

Bohgaki, M., Hakem, A., Halaby, M. J., Bohgaki, T., Li, Q., Bissey, P. A., Shloush, J., Kislinger, T., Sanchez, O., Sheng, Y., et al. (2013). The E3 ligase PIRH2 polyubiquitylates CHK2 and regulates its turnover. Cell Death Differ 20, 812–822.

Cao, C., Subhawong, T., Albert, J. M., Kim, K. W., Geng, L., Sekhar, K. R., Gi, Y. J. and Lu, B. (2006). Inhibition of mammalian target of rapamycin or apoptotic pathway induces autophagy and radiosensitizes PTEN null prostate cancer cells. Cancer Res 66, 10040–10047.

Chaurasia, M., Bhatt, A. N., Das, A., Dwarakanath, B. S. and Sharma, K. (2016). Radiation-induced autophagy: mechanisms and consequences. Free Radic Res 50, 273–290.

Chen, J. H., Zhang, P., Chen, W. D., Li, D. D., Wu, X. Q., Deng, R., Jiao, L., Li, X., Ji, J., Feng, G. K., et al. (2015). ATM-mediated PTEN phosphorylation promotes PTEN nuclear translocation and autophagy in response to DNA-damaging agents in cancer cells. Autophagy 11, 239–252.

Chen, Y., Wu, J., Liang, G., Geng, G., Zhao, F., Yin, P., Nowsheen, S., Wu, C., Li, Y., Li, L., et al. (2020). CHK2-FOXK axis promotes transcriptional control of autophagy programs. Sci Adv 6, eaax5819.

Czarny, P., Pawlowska, E., Bialkowska-Warzecha, J., Kaarniranta, K. and Blasiak, J. (2015). Autophagy in DNA damage response. Int J Mol Sci 16, 2641–2662.

Das, G., Shravage, B. V. and Baehrecke, E. H. (2012). Regulation and function of autophagy during cell survival and cell death. Cold Spring Harb Perspect Biol 4.

Denton, D. and Kumar, S. (2019). Autophagy-dependent cell death. Cell Death Differ 26, 605–616.

DeVorkin, L. and Gorski, S. M. (2014). LysoTracker staining to aid in monitoring autophagy in Drosophila. Cold Spring Harb Protoc 2014, 951–958.

Doherty, J. and Baehrecke, E. H. (2018). Life, death and autophagy. Nat Cell Biol 20, 1110–1117.

Evans, C. J., Liu, T. and Banerjee, U. (2014). Drosophila hematopoiesis: Markers and methods for molecular genetic analysis. Methods 68, 242–251.

Freeman, A. K., Dapic, V. and Monteiro, A. N. (2010). Negative regulation of CHK2 activity by protein phosphatase 2A is modulated by DNA damage. Cell Cycle 9, 736–747.

Fujiwara, K., Iwado, E., Mills, G. B., Sawaya, R., Kondo, S. and Kondo, Y. (2007). Akt inhibitor shows anticancer and radiosensitizing effects in malignant glioma cells by inducing autophagy. Int J Oncol 31, 753–760.

Gewirtz, D. A. (2014). The autophagic response to radiation: relevance for radiation sensitization in cancer therapy. Radiat Res 182, 363–367.

Golsteyn, R. M., Mundt, K. E., Fry, A. M. and Nigg, E. A. (1995). Cell cycle regulation of the activity and subcellular localization of Plk1, a human protein kinase implicated in mitotic spindle function. J Cell Biol 129, 1617–1628.

Golsteyn, R. M., Schultz, S. J., Bartek, J., Ziemiecki, A., Ried, T. and Nigg, E. A. (1994). Cell cycle analysis and chromosomal localization of human Plk1, a putative homologue of the mitotic kinases Drosophila polo and Saccharomyces cerevisiae Cdc5. J Cell Sci 107 (Pt 6), 1509–1517.

Gregory, S. L., Shandala, T., O’Keefe, L., Jones, L., Murray, M. J. and Saint, R. (2007). A Drosophila overexpression screen for modifiers of Rho signalling in cytokinesis. Fly (Austin) 1, 13–22.

Guo, Q. Q., Wang, S. S., Zhang, S. S., Xu, H. D., Li, X. M., Guan, Y., Yi, F., Zhou, T. T., Jiang, B., Bai, N., et al. (2020). ATM-CHK2-Beclin 1 axis promotes autophagy to maintain ROS homeostasis under oxidative stress. EMBO J 39, e103111.

Guo, R., Wang, S. S., Jiang, X. Y., Zhang, Y., Guo, Y., Cui, H. Y., Guo, Q. Q., Cao, L. and Xie, X. C. (2022). CHK2 Promotes Metabolic Stress-Induced Autophagy through ULK1 Phosphorylation. Antioxidants (Basel) 11.

Iliaki, S., Beyaert, R. and Afonina, I. S. (2021). Polo-like kinase 1 (PLK1) signaling in cancer and beyond. Biochem Pharmacol 193, 114747.

Inomata, K., Aoto, T., Binh, N. T., Okamoto, N., Tanimura, S., Wakayama, T., Iseki, S., Hara, E., Masunaga, T., Shimizu, H., et al. (2009). Genotoxic stress abrogates renewal of melanocyte stem cells by triggering their differentiation. Cell 137, 1088–1099.

Jang, M. S., Lee, S. J., Kim, C. J., Lee, C. W. and Kim, E. (2011). Phosphorylation by polo-like kinase 1 induces the tumor-suppressing activity of FADD. Oncogene 30, 471–481.

Khaminets, A., Ronnen-Oron, T., Baldauf, M., Meier, E. and Jasper, H. (2020). Cohesin controls intestinal stem cell identity by maintaining association of Escargot with target promoters. eLife 9, e48160.

Krzemien, J., Oyallon, J., Crozatier, M. and Vincent, A. (2010). Hematopoietic progenitors and hemocyte lineages in the Drosophila lymph gland. Dev Biol 346, 310–319.

Lécuyer, E., Parthasarathy, N. and Krause, H. M. (2008). Fluorescent In Situ Hybridization Protocols in Drosophila Embryos and Tissues. In Drosophila: Methods and Protocols (ed. C. Dahmann), pp. 289-302. Totowa, NJ: Humana Press.

Li, X., He, S. and Ma, B. (2020). Autophagy and autophagy-related proteins in cancer. Mol Cancer 19, 12.

Liu, Y., Leslie, P. L. and Zhang, Y. (2021). Life and Death Decision-Making by p53 and Implications for Cancer Immunotherapy. Trends Cancer 7, 226–239.

Ma, X., Han, Y., Song, X., Do, T., Yang, Z., Ni, J. and Xie, T. (2016). DNA damage-induced Lok/CHK2 activation compromises germline stem cell self-renewal and lineage differentiation. Development 143, 4312–4323.

Matsumura, H., Mohri, Y., Binh, N. T., Morinaga, H., Fukuda, M., Ito, M., Kurata, S., Hoeijmakers, J. and Nishimura, E. K. (2016). Hair follicle aging is driven by transepidermal elimination of stem cells via COL17A1 proteolysis. Science 351, aad4395.

Matthess, Y., Raab, M., Knecht, R., Becker, S. and Strebhardt, K. (2014). Sequential Cdk1 and Plk1 phosphorylation of caspase-8 triggers apoptotic cell death during mitosis. Mol Oncol 8, 596–608.

Moutinho-Santos, T., Sampaio, P., Amorim, I., Costa, M. and Sunkel, C. E. (1999). In vivo localisation of the mitotic POLO kinase shows a highly dynamic association with the mitotic apparatus during early embryogenesis in Drosophila. Biology of the Cell 91, 585–596.

Nguyen, T. T. N., Shim, J. and Song, Y. H. (2021). Chk2-p53 and JNK in irradiation-induced cell death of hematopoietic progenitors and differentiated cells in Drosophila larval lymph gland. Biol Open 10.

Owusu-Ansah, E. and Banerjee, U. (2009). Reactive oxygen species prime Drosophila haematopoietic progenitors for differentiation. Nature 461, 537–541.

Park, J. H., Nguyen, T. T. N., Lee, E. M., Castro-Aceituno, V., Wagle, R., Lee, K. S., Choi, J. and Song, Y. H. (2019). Role of p53 isoforms in the DNA damage response during Drosophila oogenesis. Sci Rep 9, 11473.

Pelicano, H., Carney, D. and Huang, P. (2004). ROS stress in cancer cells and therapeutic implications. Drug Resist Updat 7, 97–110.

Redza-Dutordoir, M. and Averill-Bates, D. A. (2021). Interactions between reactive oxygen species and autophagy: Special issue: Death mechanisms in cellular homeostasis. Biochim Biophys Acta Mol Cell Res 1868, 119041.

Rowe, L. A., Degtyareva, N. and Doetsch, P. W. (2008). DNA damage-induced reactive oxygen species (ROS) stress response in Saccharomyces cerevisiae. Free Radical Biology and Medicine 45, 1167–1177.

Ruf, S., Heberle, A. M., Langelaar-Makkinje, M., Gelino, S., Wilkinson, D., Gerbeth, C., Schwarz, J. J., Holzwarth, B., Warscheid, B., Meisinger, C., et al. (2017). PLK1 (polo like kinase 1) inhibits MTOR complex 1 and promotes autophagy. Autophagy 13, 486–505.

Sharma, S. K., Ghosh, S., Geetha, A. R., Mandal, S. and Mandal, L. (2019). Cell Adhesion-Mediated Actomyosin Assembly Regulates the Activity of Cubitus Interruptus for Hematopoietic Progenitor Maintenance in Drosophila. Genetics 212, 1279–1300.

Shim, J. (2015). Drosophila blood as a model system for stress sensing mechanisms. BMB Rep 48, 223–228.

Sinenko, S. A., Shim, J. and Banerjee, U. (2011). Oxidative stress in the haematopoietic niche regulates the cellular immune response in Drosophila. EMBO Rep 13, 83–89.

Smits, V. A., Klompmaker, R., Arnaud, L., Rijksen, G., Nigg, E. A. and Medema, R. H. (2000). Polo-like kinase-1 is a target of the DNA damage checkpoint. Nat Cell Biol 2, 672–676.

Tao, Y.-F., Li, Z.-H., Du, W.-W., Xu, L.-X., Ren, J.-L., Li, X.-L., Fang, F., Xie, Y., Li, M., Qian, G.-H., et al. (2017). Inhibiting PLK1 induces autophagy of acute myeloid leukemia cells via mammalian target of rapamycin pathway dephosphorylation. Oncol Rep 37, 1419–1429.

Tokusumi, T., Tokusumi, Y., Hopkins, D. W., Shoue, D. A., Corona, L. and Schulz, R. A. (2011). Germ line differentiation factor Bag of Marbles is a regulator of hematopoietic progenitor maintenance during Drosophila hematopoiesis. Development 138, 3879–3884.

Tsvetkov, L. and Stern, D. F. (2005). Phosphorylation of Plk1 at S137 and T210 is inhibited in response to DNA damage. Cell Cycle 4, 166–171.

Tsvetkov, L. M., Tsekova, R. T., Xu, X. and Stern, D. F. (2005). The Plk1 Polo box domain mediates a cell cycle and DNA damage regulated interaction with Chk2. Cell Cycle 4, 609–617.

van Vugt, M. A., Bras, A. and Medema, R. H. (2004). Polo-like kinase-1 controls recovery from a G2 DNA damage-induced arrest in mammalian cells. Mol Cell 15, 799–811.

van Vugt, M. A., Gardino, A. K., Linding, R., Ostheimer, G. J., Reinhardt, H. C., Ong, S. E., Tan, C. S., Miao, H., Keezer, S. M., Li, J., et al. (2010). A mitotic phosphorylation feedback network connects Cdk1, Plk1, 53BP1, and Chk2 to inactivate the G(2)/M DNA damage checkpoint. PLoS Biol 8, e1000287.

Wang, B., Huang, X., Liang, H., Yang, H., Guo, Z., Ai, M., Zhang, J., Khan, M., Tian, Y., Sun, Q., et al. (2021). PLK1 Inhibition Sensitizes Breast Cancer Cells to Radiation via Suppressing Autophagy. International Journal of Radiation Oncology*Biology*Physics 110, 1234–1247.

Wang, J., Sun, Q., Morita, Y., Jiang, H., Groß, A., Lechel, A., Hildner, K., Luis, Gompf, A., Hartmann, D., et al. (2012). A Differentiation Checkpoint Limits Hematopoietic Stem Cell Self-Renewal in Response to DNA Damage. Cell 148, 1001–1014.

Yoon, S., Cho, B., Shin, M., Koranteng, F., Cha, N. and Shim, J. (2017). Iron Homeostasis Controls Myeloid Blood Cell Differentiation in Drosophila. Mol Cells 40, 976–985.

Yu, L., Wan, F., Dutta, S., Welsh, S., Liu, Z., Freundt, E., Baehrecke, E. H. and Lenardo, M. (2006). Autophagic programmed cell death by selective catalase degradation. Proc Natl Acad Sci U S A 103, 4952–4957.

Yu, S., Luo, F. and Jin, L. H. (2021). Rab5 and Rab11 maintain hematopoietic homeostasis by restricting multiple signaling pathways in Drosophila. Elife 10.

Zannini, L., Delia, D. and Buscemi, G. (2014). CHK2 kinase in the DNA damage response and beyond. J Mol Cell Biol 6, 442–457.

Zhang, B., Mehrotra, S., Ng, W. L. and Calvi, B. R. (2014). Low levels of p53 protein and chromatin silencing of p53 target genes repress apoptosis in Drosophila endocycling cells. PLoS Genet 10, e1004581.

