## supplementary figures for "*polo* affects cell fate upon ionizing radiation in *Drosophila* hematopoietic progenitors by negatively regulating *lok*"

Nguyen et al. Fig S1

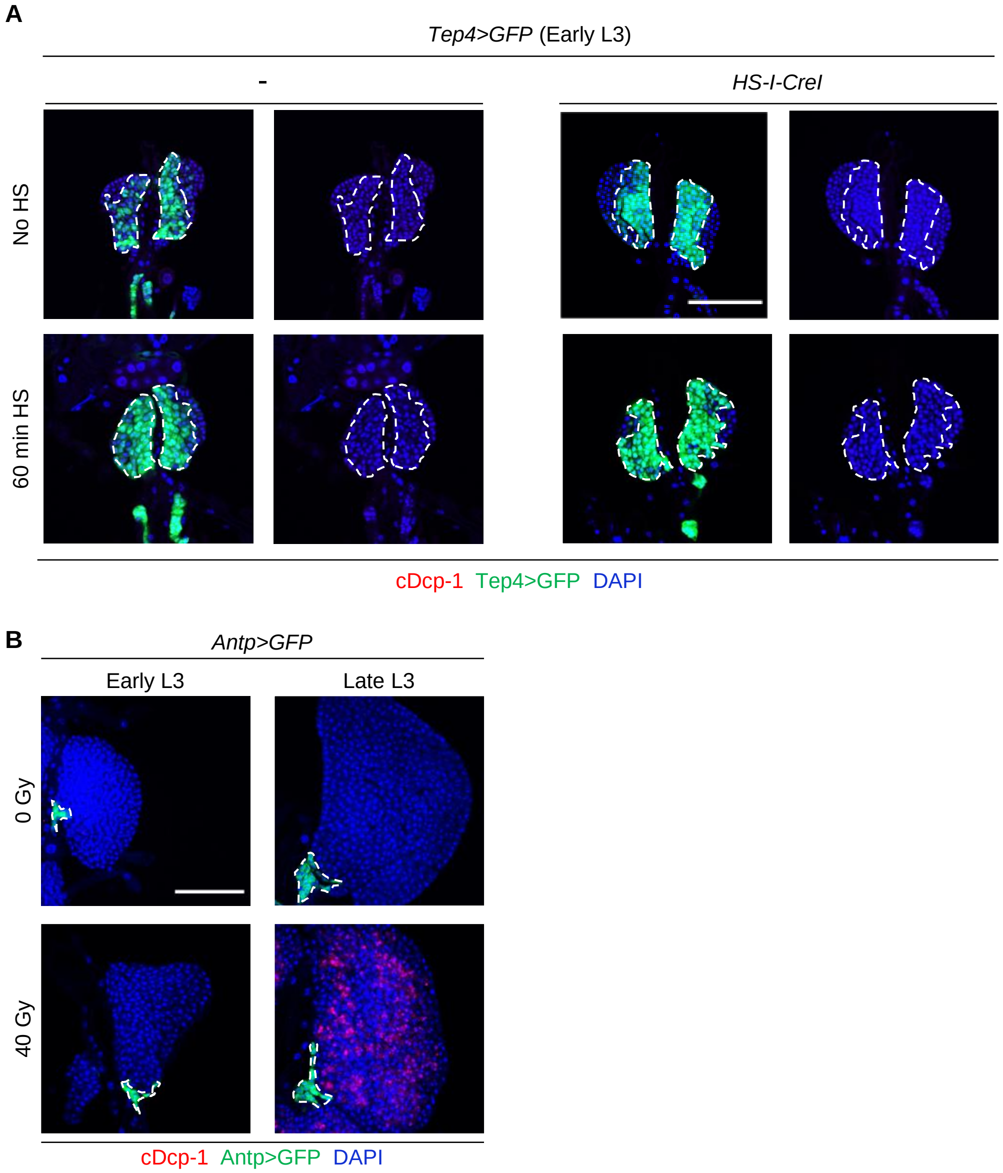


**Fig. S1. Sensitivity of lymph glands to DNA damage-induced cell death during larval development.**

(A) Early L3 larvae (44 h AEH) were heat-shock-treated to overexpress *HS-I-CreI* for 60 min. The lymph gland was stained with cDcp-1 antibody to detect apoptotic cells 4 h after treatment. Scale bars, 50 μm. DAPI (blue), Tep4>GFP (green), and cDcp-1 (red) indicate DNA, progenitors, and apoptotic cells, respectively. The boundary of the Tep4>GFP-stained MZ is marked with white broken lines. (B) Early L3 (44 h AEH) and late L3 (88 h AEH) larvae were irradiated at 40 Gy. The lymph gland was stained with cDcp-1 antibody to detect apoptotic cells 4 h after treatment. Scale bars, 50 μm. DAPI (blue), Antp>GFP (green), and cDcp-1 (red) indicate DNA, posterior signaling center (PSC) cells, and apoptotic cells, respectively. The boundary of the Antp>GFP-stained PSC cells is marked with white broken lines.

Nguyen et al. Fig S2

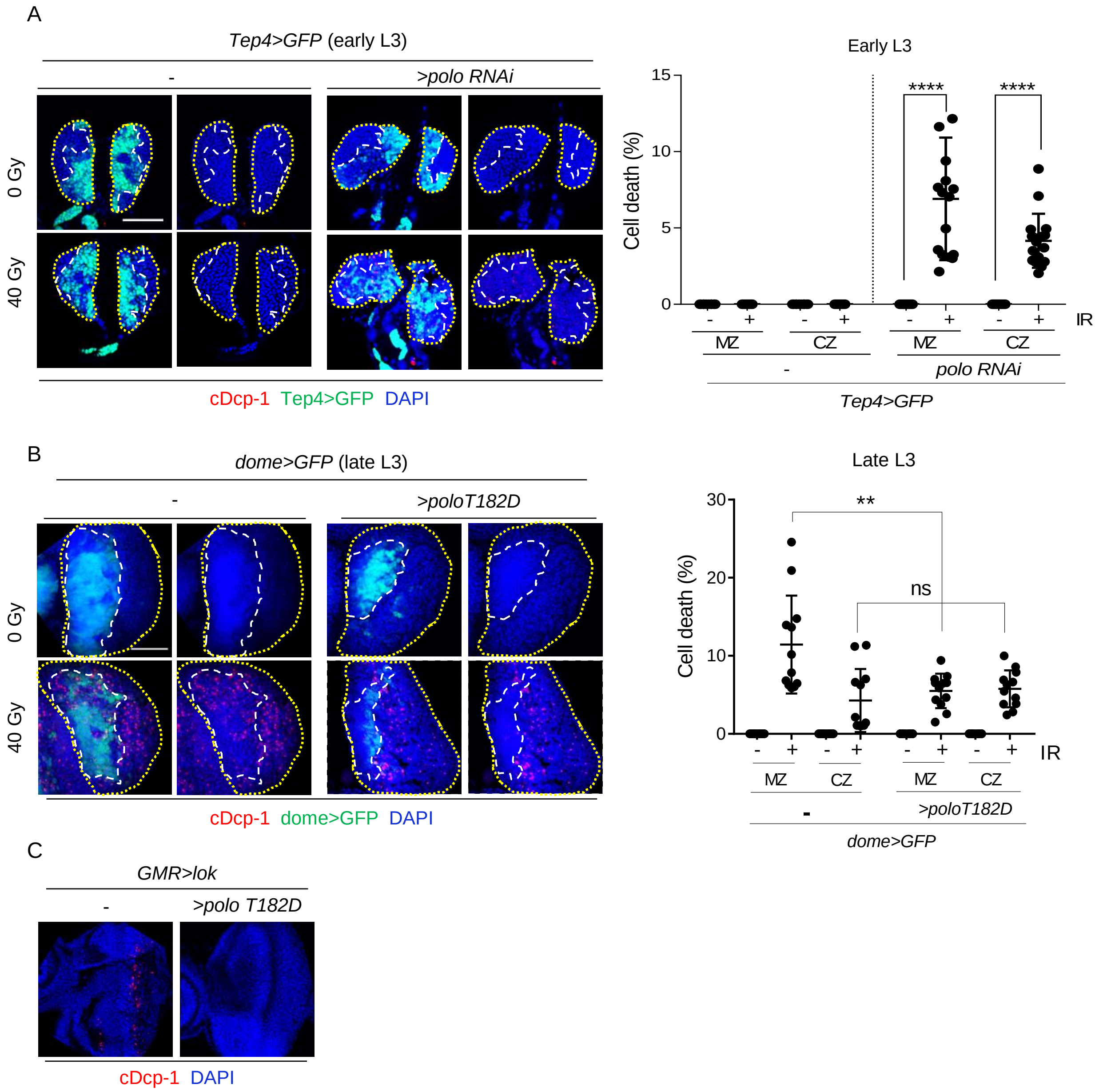


**Fig. S2. Differentially expressed *Drosophila polo* regulates cell death.**
(A, B) Early (A) or late (B) L3 expressing RNAi (*dome>polo RNAi*) (A) or the constitutively active form of *polo* (*UAS-poloT182D*) in MZ expressed by *dome-Gal4* (*dome>poloT182D*) (B) were irradiated at 40 Gy. The lymph gland was stained with cDcp-1 antibody to detect apoptotic cells 4 h after treatment. Representative images are shown. Scale bars, 50 μm. DAPI (blue), dome>GFP (green), and cDcp-1 (red) indicate DNA, progenitors, and apoptotic cells, respectively. The boundaries of the primary lobe and dome>GFP-stained MZ are marked with yellow dotted and white broken lines, respectively. The percentages of cell number with cDcp-1 signal in progenitors (MZ) and differentiated cells (CZ) are shown. Each data point represents a single primary lobe (n>10), and mean±s.d. are shown. ** *P*<0.01, **** *P*<0.0001, ns, not significant. (C) The eye disc of L3 larvae overexpressing *lok* or *lok* and *polo T182D* were stained with cDcp1. Representative images are shown.

Nguyen et al. Fig S3

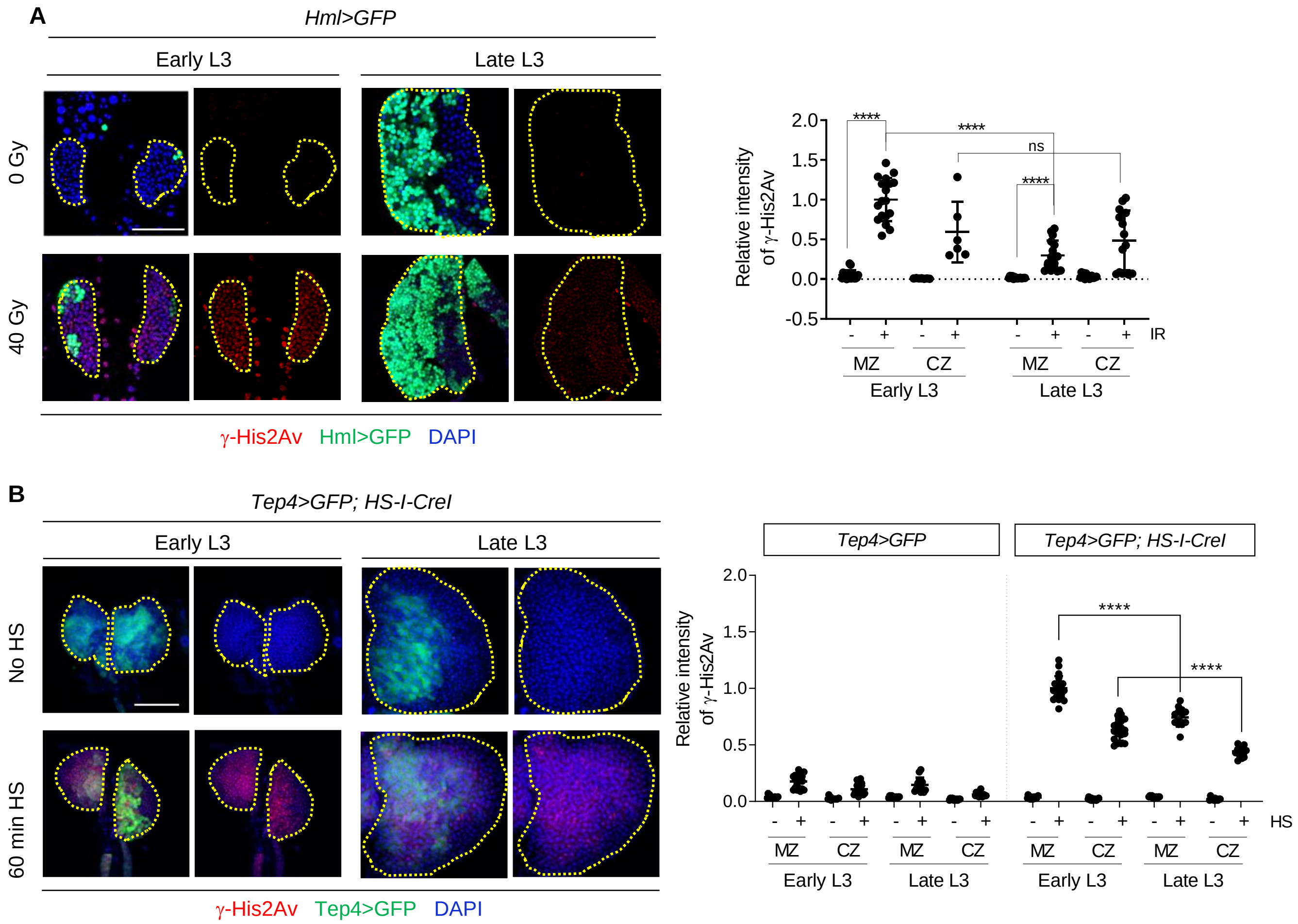


**Fig. S3. Amount of DNA damage in early and late L3 lymph glands after IR or *I-CreI* expression.**

Early and late L3 were irradiated at 40 Gy (A) or heat-shock-treated to overexpress *I-CreI* for 60 min (B). The lymph gland was stained with γ-His2Av antibody 1 h after treatment. DAPI (blue), Hml>GFP (green), Tep4>GFP (green), and γ-His2Av (red) indicate DNA, differentiated cells, progenitors, and DSBs, respectively. Scale bars, 50 μm. The boundary of the lymph gland is marked with yellow dotted lines. (A) The relative intensity of the γ-His2Av signal in progenitors (MZ) and differentiated cells (CZ) for samples in (A) with (+) and without (-) IR are shown. The values are normalized relative to the average γ-His2Av intensity in early L3 MZ after irradiation in each experiment. Each data point represents a single primary lobe (n>15), and mean±s.d. are shown. **** *P*<0.0001, ns, not significant. (B) The relative intensity of the γ-His2Av signal in progenitors (MZ) and differentiated cells (CZ) for control and L3 overexpressing *I-CreI* with (+) and without (-) heat shock are shown. The values are normalized relative to the average γ-His2Av intensity in early L3 MZ of heat-shock-treated larvae overexpressing *I-CreI* in each experiment. Each data point represents a single primary lobe (n>11), and mean±s.d. are shown. **** *P*<0.0001.

Nguyen et al. Fig S4

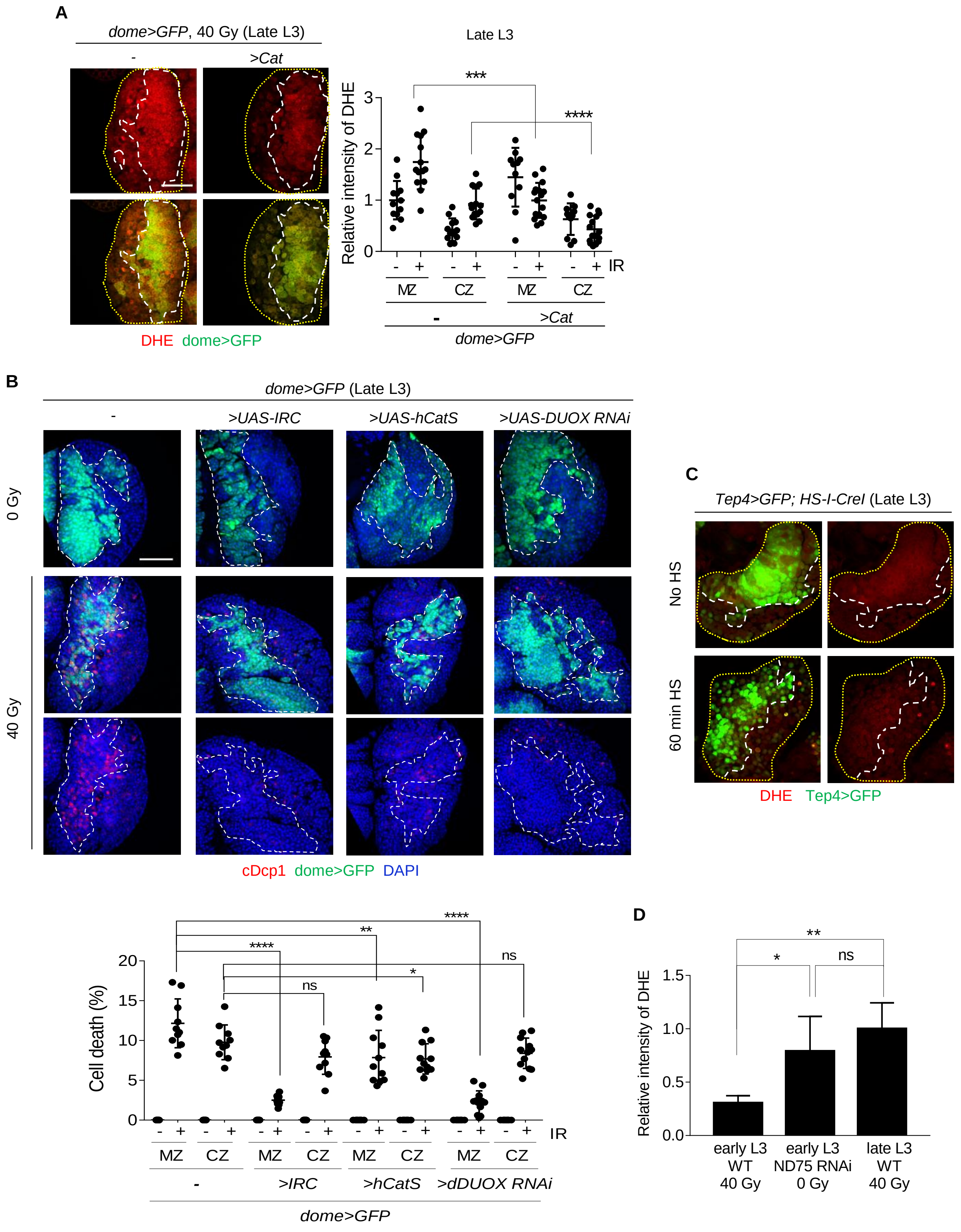

**Fig. S4. ROS induced by IR in late L3 progenitors is necessary for cell death.**

(A) Lymph glands from late L3 expressing GFP and *Catalase* in MZ (*dome>GFP, Cat*) were stained with DHE 1 h after IR. DHE (red) and *dome>GFP* (green) indicate ROS and progenitors, respectively. The boundaries of the primary lobe and MZ are marked with yellow dotted and white broken lines, respectively. Scale bars, 50 μm. The intensities of the DHE signal in progenitors (MZ) and differentiated cells (CZ) with (+) and without (-) IR are shown. The values are normalized relative to the DHE intensity in the *dome>GFP*-positive MZ in the absence of irradiation in each experiment. Each data point represents a single primary lobe (n>11), and mean±s.d. are shown. **** $P<0.0001$, *** $P<0.001$. (B) The lymph glands from late L3 expressing *IRC, hCatS,* or *Duox* RNAi in MZ were stained using cDcp-1 4 h after IR. The boundary of the MZ is marked with white broken lines. Scale bars, 50 μm. DAPI (blue), cDcp-1 (red), and dome>GFP (green) indicate DNA, apoptotic cells, and progenitors, respectively. The percentages of cDcp-1-positive cell number in progenitors (MZ) and differentiated cells (CZ) are shown. Each data point represents a single primary lobe (n=10), and mean±s.d. are shown. **** $P<0.0001$, ** $P<0.01$, * $P<0.05$, ns, not significant. (C) Late L3 were heat-shock-treated to overexpress *HS-I-CreI* for 60 min, and the lymph gland was stained with DHE 1 h after treatment. Tep4 (green) and DHE (red) indicate progenitors and ROS, respectively. The boundaries of the primary lobe and MZ are marked with yellow dotted and white broken lines, respectively. (D) The intensity of the DHE signal in irradiated early L3 that express *ND75* RNAi was compared with those in irradiated early and late L3. The values are normalized relative to the DHE intensity in the irradiated late L3 progenitors in each experiment. Each data point represents a single primary lobe (n=10), and mean±s.d. are shown. ** $P<0.01$, * $P<0.05$, ns, not significant.

Nguyen et al. Fig S5

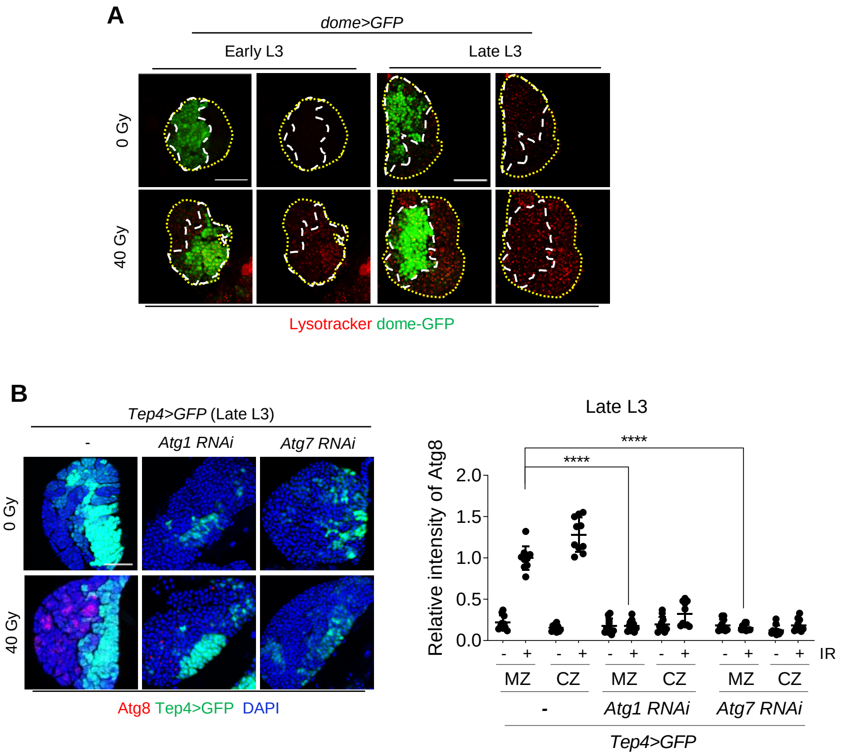


**Fig. S5. IR-induced autophagy is necessary for cell death in late L3 progenitors.**

(A) Early and late L3 were irradiated at 40 Gy. The lymph gland was stained with LysoTracker 4 h after treatment. The boundaries of the primary lobe and MZ are marked with yellow dotted and white broken lines, respectively. Scale bars, 50 μm. LysoTracker (red) and dome>GFP (green) indicate autophagy-associated lysosomal activity and progenitors, respectively. (B) Late L3 expressing RNAi against Atg1 or Atg7 using Tep4-Gal4 was irradiated at 40 Gy. The lymph glands were stained with Atg8, and the signal was quantified as described in Fig. 2. Each data point represents a single primary lobe (n=10), and mean±s.d. are shown. **** *P* <0.0001.

Nguyen et al. Fig S6

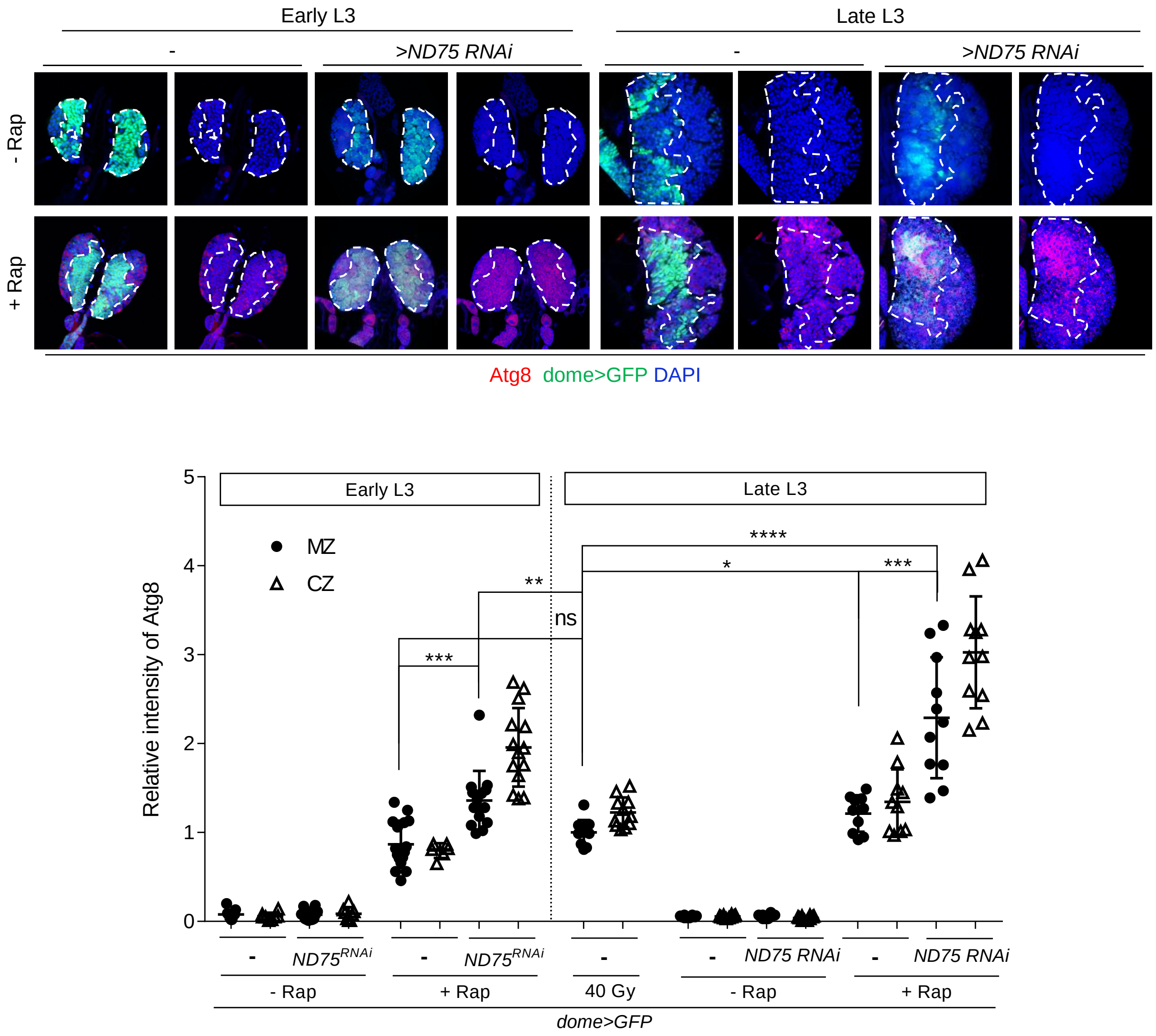


**Fig. S6. Autophagy is further enhanced by ROS induction.**
The lymph glands of early and late L3 larvae treated as described in Fig. 3 were stained with the Atg8 antibody to detect autophagosomes. DAPI (blue), Atg8 (red), and dome>GFP (green) indicate nuclei, autophagosome, and progenitor cells, respectively. Intensities of the Atg8 signal in progenitors (MZ) and differentiated cells (CZ) in (A) are shown. Each data point represents a single primary lobe (n>10), and mean±s.d. are shown. **** *P*<0.0001, *** *P*<0.001, ** *P*<0.01, * *P*<0.05, ns, not significant.

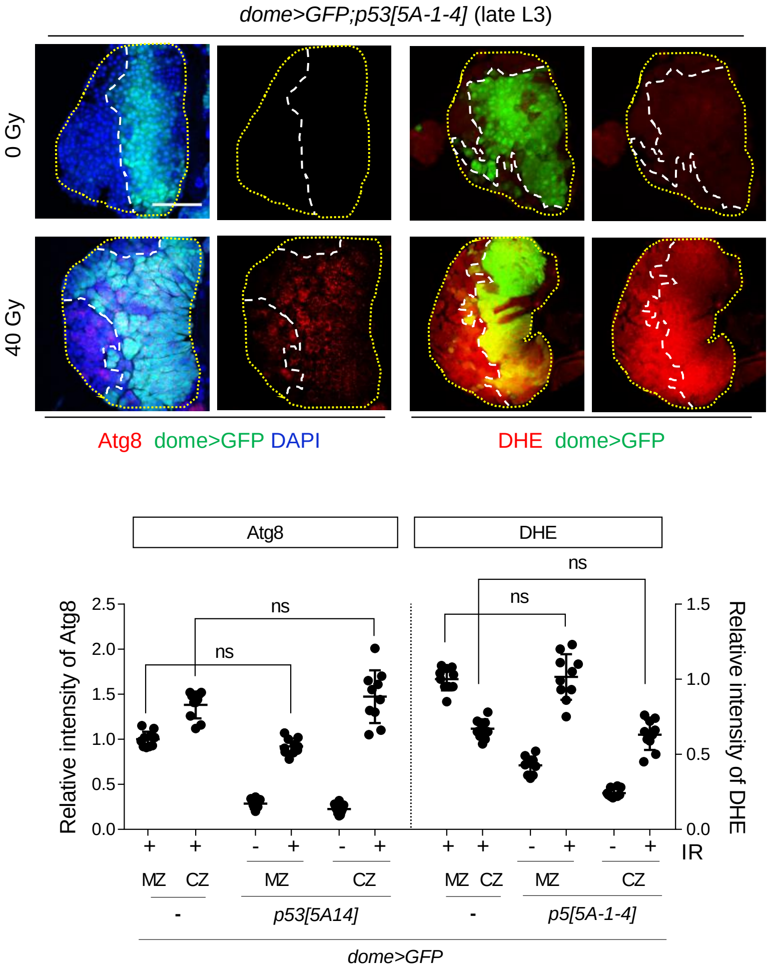


**Fig. S7. *dp53* is not required for IR-induced autophagy or ROS production in late L3 hematopoietic progenitors.**
The late L3 *p53[5A-1-4]* mutants were irradiated, and the lymph gland was stained with Atg8 antibody or DHE at 4 h or 1 h after 40 Gy irradiation, respectively. DAPI (blue), dome>GFP (green), Atg8 (red), and DHE (red) indicate DNA, progenitors, autophagosomes, and ROS, respectively. Scale bars, 50 µm. The relative intensities of Atg8 and DHE in samples were quantified. Each data point represents a single primary lobe (n=10), and mean±s.d. are shown. ns, not significant.

Nguyen et al. Fig S8

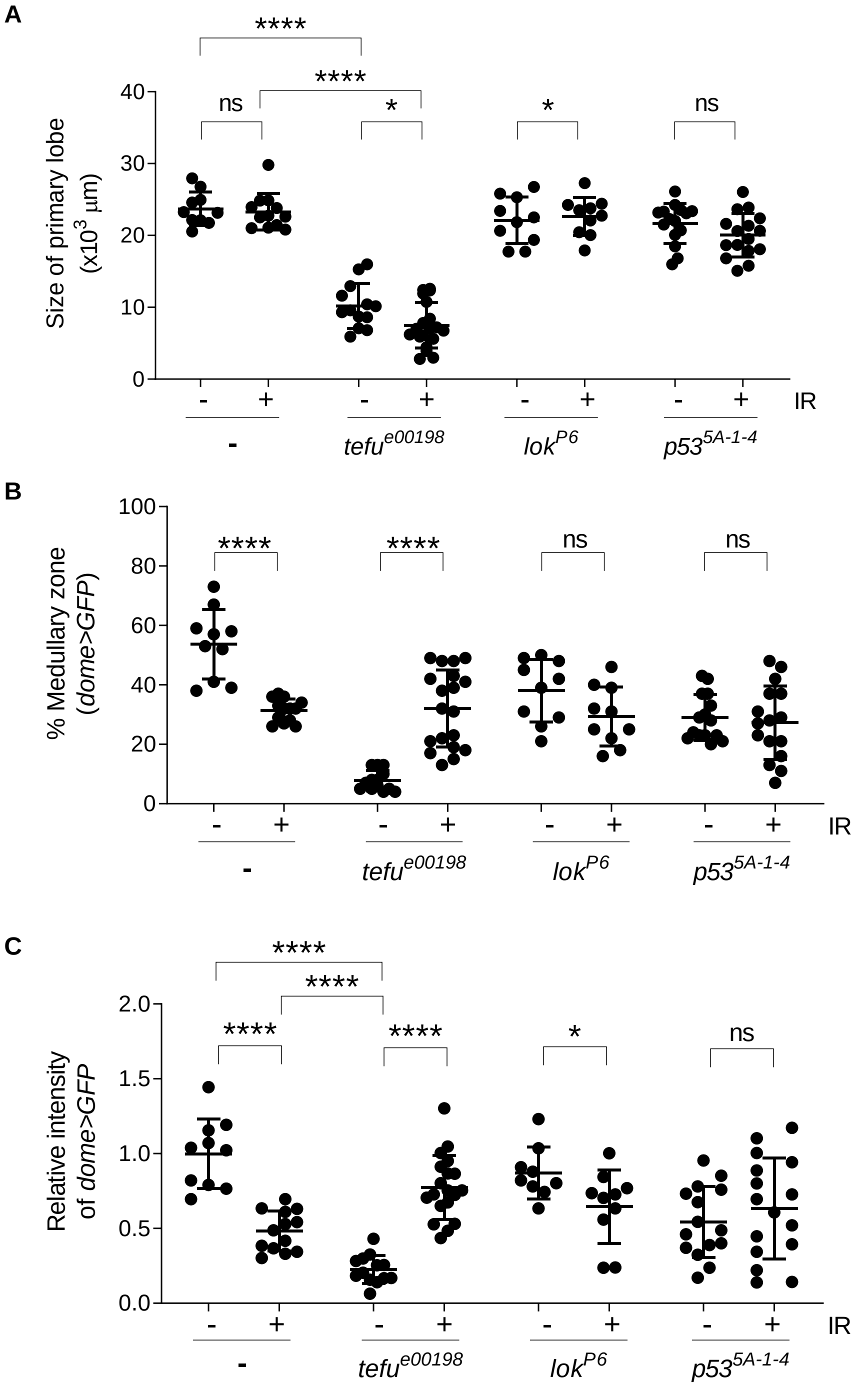


**Fig. S8. Role of DDR genes in IR-induced premature differentiation.**

For the lymph glands of irradiated early L3 in Fig. 6, the size of the primary lobes (A), percentage dome>GFP-positive area (B), and dome>GFP intensities with (+) and without (-) IR were determined. dome>GFP intensities were normalized relative to the dome>GFP intensity in the late L3 lymph gland before irradiation in each experiment. Each data point represents a single primary lobe (n>10), and mean±s.d. are shown. **** *P*<0.0001, * *P*<0.05, ns, not significant.

Nguyen et al. Fig S9

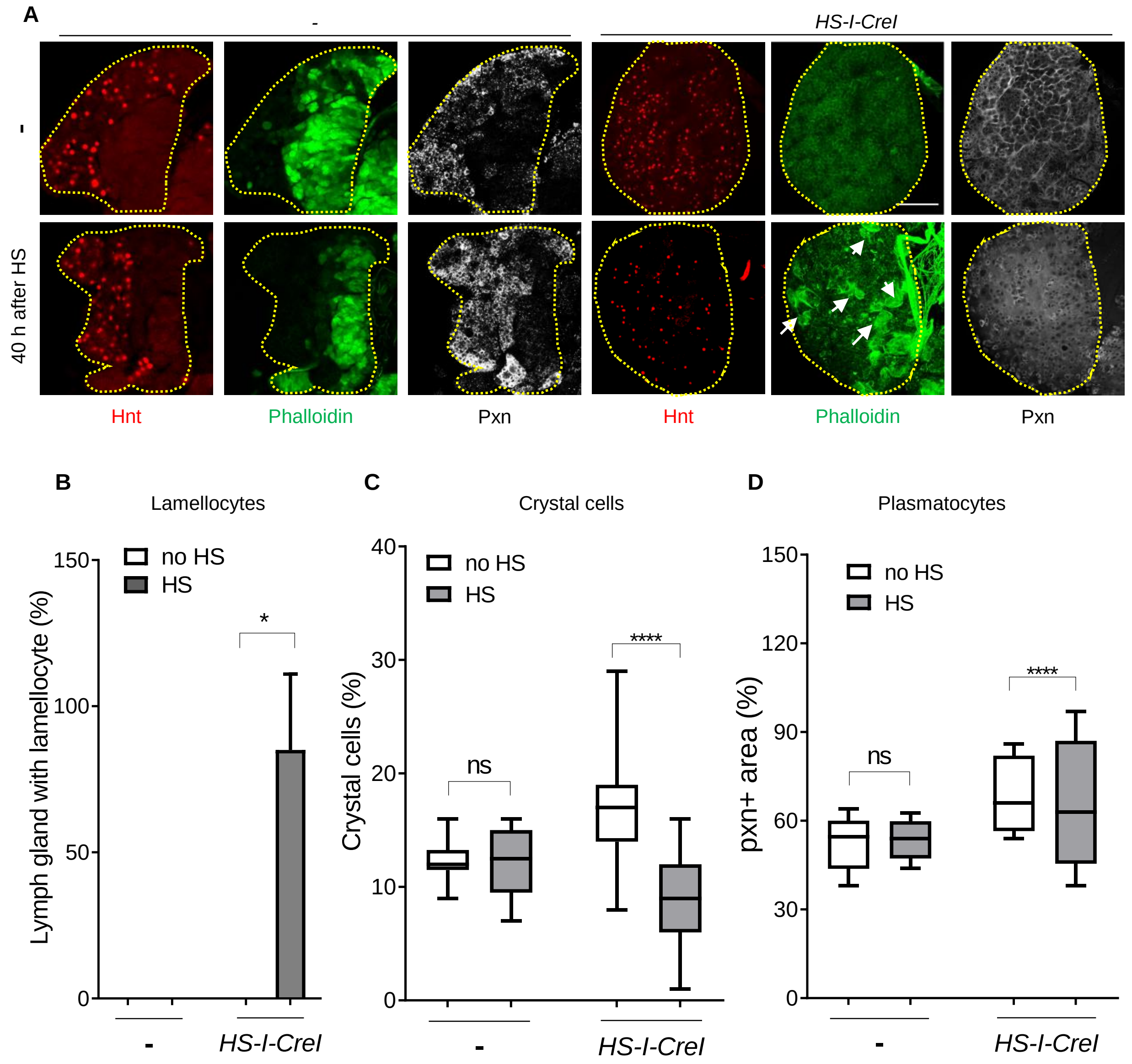


**Fig. S9. *I-CreI* expression induces premature differentiation into lamellocytes in the early L3 lymph gland.**

The early L3 were heat-shock-treated to overexpress *I-CreI* or mock-treated, and the lymph gland was stained with phalloidin, Hnt, and peroxidasin (Pxn) antibody 40 h after treatment. The boundary of the primary lobe is marked with yellow dotted lines. Scale bars, 50 µm. Phalloidin (green), Hnt (red), and Pxn (white) indicate F-actin-rich lamellocytes, crystal cells, and early stage of plasmatocytes, respectively. Arrows indicate lamellocytes. The representative images of the maximum projection of entire z-stacks of lymph glands are shown for phalloidin and Hnt. Single middle z-stack images are shown for Pxn. The lamellocytes, crystal cells, and plasmatocytes were quantified as shown in Fig. 6. Each data point represents a single primary lobe (n>10), and mean±s.d. are shown. **** *P*<0.0001, * *P*<0.05, ns, not significant.
