## Supplementary tables for "*polo* affects cell fate upon ionizing radiation in *Drosophila* hematopoietic progenitors by negatively regulating *lok*"

**Supplementary Table S1: List of genotypes and number of samples in main figures**

| **MAIN FIGURES** | | |
| --- | --- | --- |
| **Figure** | **Name of sample** | **Number of samples** |
| **Figure 1** | | |
| A | Dome-Gal4, UAS-GFP, Early L3, 0 Gy | 12 |
|  | Dome-Gal4, UAS-GFP, Early L3, 40 Gy | 16 |
|  | Dome-Gal4, UAS-GFP, Mid L3, 0 Gy | 13 |
|  | Dome-Gal4, UAS-GFP, Mid L3, 40 Gy | 11 |
| C | Tep4-Gal4, UAS-GFP/+; polo[1]/Df(rdcC-C02), Early L3, 0 Gy | 16 |
|  | Tep4-Gal4, UAS-GFP/+; polo[1]/Df(rdcC-C02), Early L3, 40 Gy | 22 |
|  | Tep4-Gal4, UAS-GFP, lok]P6]/lok[P6]; polo[1]/Df(rdcC-C02), Early L3, 0 Gy | 11 |
|  | Tep4-Gal4, UAS-GFP, lok]P6]/lok[P6]; polo[1]/Df(rdcC-C02), Early L3, 40 Gy | 10 |
| D | Tep4-Gal4, UAS-GFP, Early L3 0 Gy | 14 |
|  | Tep4-Gal4, UAS-GFP, Early L3 40 Gy 1 h | 14 |
|  | Tep4-Gal4, UAS-GFP, Early L3 40 Gy 2 h | 16 |
|  | Tep4-Gal4, UAS-GFP, Early L3 40 Gy 4 h | 14 |
|  | Tep4-Gal4, UAS-GFP, Late L3 0 Gy | 10 |
|  | Tep4-Gal4, UAS-GFP, Late L3 40 Gy 1 h | 10 |
|  | Tep4-Gal4, UAS-GFP, Late L3 40 Gy 2 h | 10 |
|  | Tep4-Gal4, UAS-GFP, Late L3 40 Gy 4 h | 12 |
| E | GFP-polo, Early L3 0 Gy | 19 |
|  | GFP-polo, Early L3 40 Gy 1 h | 25 |
|  | GFP-polo, Early L3 40 Gy 2 h | 23 |
|  | GFP-polo, Early L3 40 Gy 4 h | 17 |
|  | GFP-polo, Late L3 0 Gy | 15 |
|  | GFP-polo, Late L3 40 Gy 1 h | 16 |
|  | GFP-polo, Late L3 40 Gy 2 h | 18 |
|  | GFP-polo, Late L3 40 Gy 4 h | 11 |
| F | Tep4-Gal4, UAS-GFP, Early L3 0 Gy | 12 |
|  | Tep4-Gal4, UAS-GFP, Early L3 40 Gy 1 h | 14 |
|  | Tep4-Gal4, UAS-GFP, Early L3 40 Gy 2 h | 16 |
|  | Tep4-Gal4, UAS-GFP, Early L3 40 Gy 4 h | 12 |
|  | Tep4-Gal4, UAS-GFP, Late L3 0 Gy | 10 |
|  | Tep4-Gal4, UAS-GFP, Late L3 40 Gy 1 h | 10 |
|  | Tep4-Gal4, UAS-GFP, Late L3 40 Gy 2 h | 10 |
|  | Tep4-Gal4, UAS-GFP, Late L3 40 Gy 4 h | 10 |
| **Figure 2** | | |
| A | Dome-Gal4, UAS-GFP, Early L3, 0 Gy | 31 |
|  | Dome-Gal4, UAS-GFP, Early L3, 40 Gy | 23 |
|  | Dome-Gal4, UAS-GFP, Late L3, 0 Gy | 23 |
|  | Dome-Gal4, UAS-GFP, Late L3, 40 Gy | 18 |
| B | Dome-Gal4, UAS-GFP, Late L3, 0 Gy | 12 |
|  | Dome-Gal4, UAS-GFP, Late L3, 40 Gy | 19 |
|  | Dome-Gal4, UAS-GFP/+; UAS-Cat/+ Late L3, 0 Gy | 15 |
|  | Dome-Gal4, UAS-GFP/+; UAS-Cat/+, Late L3, 40 Gy | 15 |
| C | Tep4-Gal4, UAS-GFP, Early 3L 0 Gy | 11 |
|  | Tep4-Gal4, UAS-GFP, Early 3L 40 Gy | 10 |
|  | Tep4-Gal4, UAS-GFP, Late 3L 0 Gy | 10 |
|  | Tep4-Gal4, UAS-GFP, Late 3L 40 Gy | 12 |
| D | Tep4-Gal4, UAS-GFP, Late 3L 0 Gy | 10 |
|  | Tep4-Gal4, UAS-GFP, Late 3L 40 Gy | 10 |
|  | Tep4-Gal4, UAS-GFP, Late 3L, CQ | 10 |
|  | Tep4-Gal4, UAS-GFP, Late 3L, 40 Gy, CQ | 10 |
| E | Tep4-Gal4, UAS-GFP, Late 3L 0 Gy | 10 |
|  | Tep4-Gal4, UAS-GFP, Late 3L 40 Gy | 10 |
|  | Tep4-Gal4, UAS-GFP/Atg1 RNAi, Late 3L 0 Gy | 10 |
|  | Tep4-Gal4, UAS-GFP/Atg1 RNAi, Late 3L 40 Gy | 10 |
|  | Tep4-Gal4, UAS-GFP/+; Atg7 RNAi/+, Late 3L 0 Gy | 10 |
|  | Tep4-Gal4, UAS-GFP/+; Atg7 RNAi/+, Late 3L 40 Gy | 10 |
| F | Tep4-Gal4, UAS-GFP, Late 3L 0 Gy | 11 |
|  | Tep4-Gal4, UAS-GFP, Late 3L 40 Gy | 11 |
|  | Tep4-Gal4, UAS-GFP/Atg1 RNAi, Late 3L 0 Gy | 10 |
|  | Tep4-Gal4, UAS-GFP/Atg1 RNAi, Late 3L 40 Gy | 10 |
|  | Tep4-Gal4, UAS-GFP/+; Atg7 RNAi/+, Late 3L 0 Gy | 10 |
|  | Tep4-Gal4, UAS-GFP/+; Atg7 RNAi/+, Late 3L 40 Gy | 11 |
| G | Dome-Gal4, UAS-GFP, Late L3, 0 Gy | 11 |
|  | Dome-Gal4, UAS-GFP, Late L3, 40 Gy | 11 |
|  | Dome-Gal4, UAS-GFP/+; UAS-Cat/+ Late L3, 0 Gy | 11 |
|  | Dome-Gal4, UAS-GFP/+; UAS-Cat/+, Late L3, 40 Gy | 12 |
| **Figure 3** | | |
|  | Dome-Gal4, UAS-GFP, Early L3, no Rap | 28 |
|  | Dome-Gal4, UAS-GFP/+; ND75 RNAi/+, Early L3, no Rap | 24 |
|  | Dome-Gal4, UAS-GFP, Early L3, Rap | 10 |
|  | Dome-Gal4, UAS-GFP/+; ND75 RNAi/+, Early L3, Rap | 14 |
|  | Dome-Gal4, UAS-GFP, Early L3, 40 Gy | 17 |
|  | Dome-Gal4, UAS-GFP/+; ND75 RNAi/+, Early L3, 40 Gy | 18 |
|  | Dome-Gal4, UAS-GFP, Late L3, no Rap | 21 |
|  | Dome-Gal4, UAS-GFP/+; ND75 RNAi/+, Late L3, no Rap | 27 |
|  | Dome-Gal4, UAS-GFP, Late L3, Rap | 10 |
|  | Dome-Gal4, UAS-GFP/+; ND75 RNAi/+, Late L3, Rap | 10 |
| **Figure 4** | | |
| A | Atg8, Tep4-Gal4, UAS-GFP/+; polo[1]/Df(rdcC-C02), Early L3, 0 Gy | 24 |
|  | Atg8, Tep4-Gal4, UAS-GFP/+; polo[1]/Df(rdcC-C02), Early L3, 40 Gy | 34 |
|  | DHE, Tep4-Gal4, UAS-GFP, lok]P6]/lok[P6]; polo[1]/Df(rdcC-C02), Early L3, 0 Gy | 12 |
|  | DHE, Tep4-Gal4, UAS-GFP, lok]P6]/lok[P6]; polo[1]/Df(rdcC-C02), Early L3, 40 Gy | 10 |
|  | DHE, Tep4-Gal4, UAS-GFP/+; polo[1]/Df(rdcC-C02), Early L3, 0 Gy | 16 |
|  | DHE, Tep4-Gal4, UAS-GFP/+; polo[1]/Df(rdcC-C02), Early L3, 40 Gy | 13 |
|  | DHE, Tep4-Gal4, UAS-GFP, lok]P6]/lok[P6]; polo[1]/Df(rdcC-C02), Early L3, 0 Gy | 12 |
|  | DHE, Tep4-Gal4, UAS-GFP, lok]P6]/lok[P6]; polo[1]/Df(rdcC-C02), Early L3, 40 Gy | 13 |
| D | Dome-Gal4, UAS-GFP, Late L3, 40 Gy | 16 |
|  | Dome-Gal4, UAS-GFP/+; UAS-lok/+, Early L3 | 10 |
|  | Dome-Gal4, UAS-GFP/+; UAS-lok/+, Late L3 | 10 |
|  | Dome-Gal4, UAS-GFP/+; UAS-poloT182D/+, Late L3, 0 Gy | 10 |
|  | Dome-Gal4, UAS-GFP/+; UAS-poloT182D/+, Late L3, 40 Gy | 12 |
|  | Dome-Gal4, UAS-GFP/+; lok]P6]/lok[P6], Late L3, 0 Gy | 10 |
|  | Dome-Gal4, UAS-GFP/+; lok]P6]/lok[P6]; Late L3, 40 Gy | 10 |
| E | Dome-Gal4, UAS-GFP, Late L3, 40 Gy | 20 |
|  | Dome-Gal4, UAS-GFP/+; UAS-lok/+, Early L3 | 11 |
|  | Dome-Gal4, UAS-GFP/+; UAS-lok/+, Late L3 | 10 |
|  | Dome-Gal4, UAS-GFP/+; UAS-poloT182D/+, Late L3, 0 Gy | 10 |
|  | Dome-Gal4, UAS-GFP/+; UAS-poloT182D/+, Late L3, 40 Gy | 10 |
|  | Dome-Gal4, UAS-GFP/+; lok]P6]/lok[P6], Late L3, 0 Gy | 11 |
|  | Dome-Gal4, UAS-GFP/+; lok]P6]/lok[P6]; Late L3, 40 Gy | 11 |
| **Figure 5** | | |
|  | Dome-Gal4, UAS-GFP, Early L3, 0 Gy | 31 |
|  | Dome-Gal4, UAS-GFP, Early L3, 40 Gy | 28 |
| **Figure 6** | | |
| B | CS, 0 Gy | 15 |
|  | CS, 40 Gy | 13 |
|  | tefu[e0198], 0 Gy | 10 |
|  | tefu[e0198], 40 Gy | 17 |
|  | lok[P6], 0 Gy | 14 |
|  | lok [P6], 40 Gy | 15 |
|  | p53[5A14], 0 Gy | 14 |
|  | p53[5A14], 40 Gy | 15 |
| C | CS, 0 Gy | 15 |
|  | CS, 40 Gy | 13 |
|  | tefu[e0198], 0 Gy | 13 |
|  | tefu[e0198], 40 Gy | 18 |
|  | lok[P6], 0 Gy | 22 |
|  | lok [P6], 40 Gy | 18 |
|  | p53[5A14], 0 Gy | 11 |
|  | p53[5A14], 40 Gy | 17 |
| D | CS, 0 Gy | 15 |
|  | CS, 40 Gy | 13 |
|  | tefu[e0198], 0 Gy | 10 |
|  | tefu[e0198], 40 Gy | 17 |
|  | lok[P6], 0 Gy | 14 |
|  | lok [P6], 40 Gy | 15 |
|  | p53[5A14], 0 Gy | 14 |
|  | p53[5A14], 40 Gy | 15 |

**Supplementary Table S2: List of genotypes and number of samples in supplementary figures**

| **SUPPLEMENTARY FIGURES** | | |
| --- | --- | --- |
| **Figure** | **Name of sample** | **Number of samples** |
| **Figure S2** | | |
| A | Tep4-Gal4, UAS-GFP, Early L3 0 Gy | 10 |
|  | Tep4-Gal4, UAS-GFP, Early L3 40 Gy | 10 |
|  | Tep4-Gal4, UAS-GFP/+; polo RNAi, Early L3 0 Gy | 11 |
|  | Tep4-Gal4, UAS-GFP/+; polo RNAi, Early L3 40 Gy | 16 |
| B | Dome-Gal4, UAS-GFP, Late L3, 0 Gy | 12 |
|  | Dome-Gal4, UAS-GFP, Late L3, 40 Gy | 12 |
|  | Dome-Gal4, UAS-GFP/+; UAS-poloT182D/+, Late L3, 0 Gy | 10 |
|  | Dome-Gal4, UAS-GFP/+; UAS-poloT182D/+, Late L3, 40 Gy | 12 |
| **Figure S3** | | |
| A | Hml-Gal4, UAS-GFP, Early L3, 0 Gy | 17 |
|  | Hml-Gal4, UAS-GFP, Early L3, 40 Gy | 18 |
|  | Hml-Gal4, UAS-GFP, Late L3, 0 Gy | 15 |
|  | Hml-Gal4, UAS-GFP, Late L3, 40 Gy | 16 |
| B | Tep4-Gal4, UAS-GFP, Early L3, no heat shock | 15 |
|  | Tep4-Gal4, UAS-GFP, Early L3, heat shock | 20 |
|  | Tep4-Gal4, UAS-GFP, Late 3L, no heat shock | 11 |
|  | Tep4-Gal4, UAS-GFP, Late 3L, heat shock | 14 |
|  | Tep4-Gal4, UAS-GFP/+; HS-ICreI/+, Early L3, no heat shock | 16 |
|  | Tep4-Gal4, UAS-GFP/+; HS-ICreI/+, Early L3, heat shock | 19 |
|  | Tep4-Gal4, UAS-GFP/+; HS-ICreI/+, Late 3L, no heat shock | 11 |
|  | Tep4-Gal4, UAS-GFP/+; HS-ICreI/+, Late 3L, heat shock | 14 |
| **Figure S4** | | |
| A | Dome-Gal4, UAS-GFP, Late L3, 0 Gy | 12 |
|  | Dome-Gal4, UAS-GFP, Late L3, 40 Gy | 14 |
|  | Dome-Gal4, UAS-GFP/+; UAS-Cat/+ Late L3, 0 Gy | 11 |
|  | Dome-Gal4, UAS-GFP/+; UAS-Cat/+, Late L3, 40 Gy | 16 |
| B | Dome-Gal4, UAS-GFP, Late L3, 0 Gy | 11 |
|  | Dome-Gal4, UAS-GFP, Late L3, 40 Gy | 10 |
|  | Dome-Gal4, UAS-GFP/+; UAS-IRC/+, Late L3, 0 Gy | 11 |
|  | Dome-Gal4, UAS-GFP/+; UAS-IRC/+, Late L3, 40 Gy | 10 |
|  | Dome-Gal4, UAS-GFP/+; UAS-hCatS/+, Late L3, 0 Gy | 10 |
|  | Dome-Gal4, UAS-GFP/+; UAS-hCatS/+, Late L3, 40 Gy | 11 |
|  | Dome-Gal4, UAS-GFP/+; dDUOX RNAi/+, Late L3, 0 Gy | 10 |
|  | Dome-Gal4, UAS-GFP/+; dDUOX RNAi/+, Late L3, 40 Gy | 12 |
| D | Dome-Gal4, UAS-GFP, Early L3, 40 Gy | 10 |
|  | Dome-Gal4, UAS-GFP/+; ND75 RNAi/+, Early L3, no Rap | 10 |
|  | Dome-Gal4, UAS-GFP, Late L3, 40 Gy | 10 |
| **Figure S5** | | |
| B | Tep4-Gal4, UAS-GFP, Late 3L 0 Gy | 10 |
|  | Tep4-Gal4, UAS-GFP, Late 3L 40 Gy | 10 |
|  | Tep4-Gal4, UAS-GFP/Atg1 RNAi, Late 3L 0 Gy | 10 |
|  | Tep4-Gal4, UAS-GFP/Atg1 RNAi, Late 3L 40 Gy | 10 |
|  | Tep4-Gal4, UAS-GFP/+; Atg7 RNAi/+, Late 3L 0 Gy | 10 |
|  | Tep4-Gal4, UAS-GFP/+; Atg7 RNAi/+, Late 3L 40 Gy | 10 |
| **Figure S6** | | |
|  | Dome-Gal4, UAS-GFP, Early L3, no Rap | 11 |
|  | Dome-Gal4, UAS-GFP/+; ND75 RNAi/+, Early L3, no Rap | 14 |
|  | Dome-Gal4, UAS-GFP, Early L3, Rap | 16 |
|  | Dome-Gal4, UAS-GFP/+; ND75 RNAi/+, Early L3, Rap | 14 |
|  | Dome-Gal4, UAS-GFP, Late L3, 40 Gy | 11 |
|  | Dome-Gal4, UAS-GFP, Late L3, no Rap | 10 |
|  | Dome-Gal4, UAS-GFP/+; ND75 RNAi/+, Late L3, no Rap | 10 |
|  | Dome-Gal4, UAS-GFP, Late L3, Rap | 10 |
|  | Dome-Gal4, UAS-GFP/+; ND75 RNAi/+, Late L3, Rap | 11 |
| **Figure S7** | | |
|  | Atg8, Dome-Gal4, UAS-GFP, Late L3, 0 Gy | 10 |
|  | Atg8, Dome-Gal4, UAS-GFP, Late L3, 40 Gy | 10 |
|  | Atg8, Dome-Gal4, UAS-GFP, p53[5A-1-4], Late L3, 0 Gy | 10 |
|  | Atg8, Dome-Gal4, UAS-GFP, p53[5A-1-4], Late L3, 40 Gy | 10 |
|  | DHE, Dome-Gal4, UAS-GFP, Late L3, 0 Gy | 10 |
|  | DHE, Dome-Gal4, UAS-GFP, Late L3, 40 Gy | 10 |
|  | DHE, Dome-Gal4, UAS-GFP, p53[5A-1-4], Late L3, 0 Gy | 10 |
|  | DHE, Dome-Gal4, UAS-GFP, p53[5A-1-4], Late L3, 40 Gy | 10 |
| **Figure S8** | | |
| A, B, C | Dome-Gal4, UAS-GFP; Hml-dsRed, 0 Gy | 10 |
|  | Dome-Gal4, UAS-GFP; Hml-dsRed, 40 Gy | 12 |
|  | Dome-Gal4, UAS-GFP; e00198, 0 Gy | 13 |
|  | Dome-Gal4, UAS-GFP; e00198, 40 Gy | 19 |
|  | Dome-Gal4, UAS-GFP; lok[P6], 0 Gy | 10 |
|  | Dome-Gal4, UAS-GFP; lok[P6], 40 Gy | 10 |
|  | Dome-Gal4, UAS-GFP, p53[5A-1-4], 0 Gy | 15 |
|  | Dome-Gal4, UAS-GFP, p53[5A-1-4], 40 Gy | 16 |
| **Figure S9** | | |
| B, C, D | Tep4-Gal4, UAS-GFP, no heat shock | 10 |
|  | Tep4-Gal4, UAS-GFP, heat shock | 10 |
|  | Tep4-Gal4, UAS-GFP/+; HS-ICreI/+, no heat shock | 17 |
|  | Tep4-Gal4, UAS-GFP/+; HS-ICreI/+, heat shock | 21 |
